# Altered immunity of laboratory mice in the natural environment is associated with fungal colonization

**DOI:** 10.1101/742387

**Authors:** Frank Yeung, Ying-Han Chen, Jian-Da Lin, Jacqueline M. Leung, Caroline McCauley, Joseph C. Devlin, Christina Hansen, Alex Cronkite, Zac Stephens, Charlotte Drake-Dunn, Yi Fulmer, Bo Shopsin, Kelly V. Ruggles, June L. Round, P’ng Loke, Andrea L. Graham, Ken Cadwell

**Author notes:** These authors contributed equally (Frank Yeung, Ying-Han Chen, Jian-Da Lin). These authors contributed equally (P’ng Loke, Andrea L. Graham, Ken Cadwell).

## Abstract

The immune systems of free-living mammals such as humans and wild mice display a heightened degree of activation compared with laboratory mice maintained under artificial conditions. Here, we demonstrate that releasing inbred laboratory mice into an outdoor enclosure to mimic life in a natural environment alters the state of immunity. In addition to enhancing the differentiation of T cell populations previously associated with pathogen exposure, we found that outdoor release of mice led to an increase in circulating granulocytes. However, rewilded mice were not infected by pathogens previously implicated in immune activation. Rather, changes to the immune system were associated with an altered composition of the microbiota, and fungi isolated from rewilded mice were sufficient to increase circulating granulocytes. These findings establish an experimental procedure to investigate the impact of the natural environment on immune development and identify a role for sustained fungal exposure in determining granulocyte numbers.

## Introduction

The house mouse is the most prevalent mammalian model organism used in biomedical research and has enabled fundamental advances in basic immunology. Yet, this ubiquitous model fails to recreate certain aspects of human immunity. Inbred laboratory mice and adult humans differ in the proportion of leukocyte subsets, transcriptional responses to microbial challenges, and other immune parameters (Masopust et al., 2017; Tao and Reese, 2017). Such differences may limit the predictive value of experiments with mice when studying complex inflammatory and infectious diseases, resulting in significant shortcomings in translating laboratory observations to humans.

Recent findings suggest that this shortcoming of the rodent model may be due to the specific pathogen free (SPF) environment in which they are maintained. Wild mice and pet store mice, both of which are exposed to a litany of pathogens that are typically excluded from SPF facilities, display an abundance of differentiated memory T cells that more closely resembles the state of immunity in adult humans (Abolins et al., 2017; Beura et al., 2016; Choi et al., 2019). Similarly, transferring embryos from lab mice into wild mice generates commensal- and pathogen-exposed offspring (wildlings) that more faithfully recreate human immunity than standard SPF mice, including the unresponsiveness to immunotherapies that failed in clinical trials (Rosshart et al., 2019). Sequentially infecting SPF mice with 3 viruses and a helminth shifts the gene expression profile of peripheral blood mononuclear cells (PBMCs) towards that of pet store mice and adult humans (Reese et al., 2016), further highlighting the role for pathogen experience in normalizing the immune system. SPF mice are also distinguished from free-living mammals by the lack of exposure to potentially immuno-stimulatory members of the microbiota that are absent in a laboratory animal facility. For example, the offspring of germ-free mice inoculated with ileocecal contents from wild mice display increased resistance to influenza infection and colorectal tumorigenesis (Rosshart et al., 2017). However, the specific effect of the naturally-acquired microbiota on immune development is unclear, and the non-bacterial members of the wild microbiota such as fungi have not been examined in detail.

We recently described a mesocosm system in which SPF mice are ‘rewilded’ through controlled release into an outdoor enclosure facility (Leung et al., 2018). In contrast to approaches that rely on catching wild mice, our experimental system allows us to control genetics and the timing of environmental exposure by releasing lab-bred mice into a natural environment. We can thus systematically compare genetically defined mice maintained outdoors with control mice of the same genotype kept in a conventional animal facility. Another key feature of the enclosure is that a zinced iron wall excludes predators and rodents harboring disease-causing infectious agents, while allowing exposure to natural soil, vegetation, and weather. Mice rewilded through transient release into the enclosure acquire a bacterial microbiota characterized by increased diversity and display heightened susceptibility to helminth infection (Leung et al., 2018).

In a companion manuscript (Lin *et al*), we describe deep immune-profiling of rewilded mice to identify a series of changes that occur to the immune system upon exposure to the natural environment, and apply computational analyses to infer the impact of genetic and environmental variables. Here, we tested the hypothesis that colonization by microorganisms present in the natural environment contribute to the maturation of the immune system that occurs in free-living mammals. We found that rewilded mice display an increase in central and effector T cells similar to pet store mice infected with pathogens. We also identified a new feature, an increase in granulocyte populations in the blood and lymph node. These changes were not associated with pathogen infection, and instead, were associated with an altered gut microbiota characterized by a substantial increase in fungi. Colonization of lab mice with the model fungus *Candida albicans* or fungi isolated from rewilded mice recreated the expansion of granulocytes. These findings implicate fungi and other members of the gut microbiota in the maturation of the immune system, and suggest that differences between lab mice and humans are due, in part, to lack of colonization by environmental microbes.

## RESULTS

### Rewilding increases the maturation of lymphocytes

To determine the consequences of microbial colonization and exposure to the natural environment on the steady-state immune system, we applied multicolor flow cytometry to analyze the immune cell composition of blood and mesenteric lymph nodes (MLNs) from SPF mice aged 6-8 weeks old at the time of release into the enclosure and captured 6-7 weeks later. Details of the cohort of rewilded mice are described in the Methods and includes male and female wild-type C57BL6/J mice, and also an additional cohort of mice harboring mutations in inflammatory bowel disease (IBD) genes (*Atg16L1* and *Nod2*) described in the companion study (Lin *et. al.*). Matched control mice were maintained under SPF conditions in the institutional vivarium, herein referred to as lab mice. The initial flow cytometry analyses were performed on all rewilded and lab mice as genotype did not affect the outcome (Lin *et. al.*). We provide the detailed breakdown by *Atg16L1* and *Nod2* mutation status in subsequent analyses.

In contrast to adult humans and pet store mice, SPF mice and human neonates lack differentiated memory CD8^+^ T cells harboring CD44 and other markers of antigen experience (Beura et al., 2016). We found that release into the outdoor enclosure led to the acquisition of these signatures of differentiation in T cells from the blood and MLNs. Compared with lab control mice, rewilded mice had CD8^+^ T cells with surface markers identifying them as CD62L^lo^CD44^hi^ effector memory (T_EM_) and CD62L^hi^CD44^hi^ central memory (T_CM_) populations, and a corresponding decrease in CD62L^hi^CD44^lo^ naïve T cells (Fig. 1A to B). Similar increases in differentiated CD4^+^ T cell populations were observed in rewilded mice (fig. S1, A to B). We also found an increase in CD8^+^ and CD4^+^ T cells expressing activation markers KLRG1 and CD25, respectively (Fig. 1C, and fig. S2C). Another striking feature of pet shop mice is the increase in mucosally-distributed T cells compared with lab mice. Histological examination of the small intestine from rewilded mice revealed an increased number of CD8^+^ cells (Fig. 1D). Despite these features of enhanced immune activation, we did not detect inflammatory lesions in intestinal sections (Fig. 1D). Thus, rewilding is associated with a general enhancement in T cell maturation resembling observations made in pet store mice.

**Figure 1.**
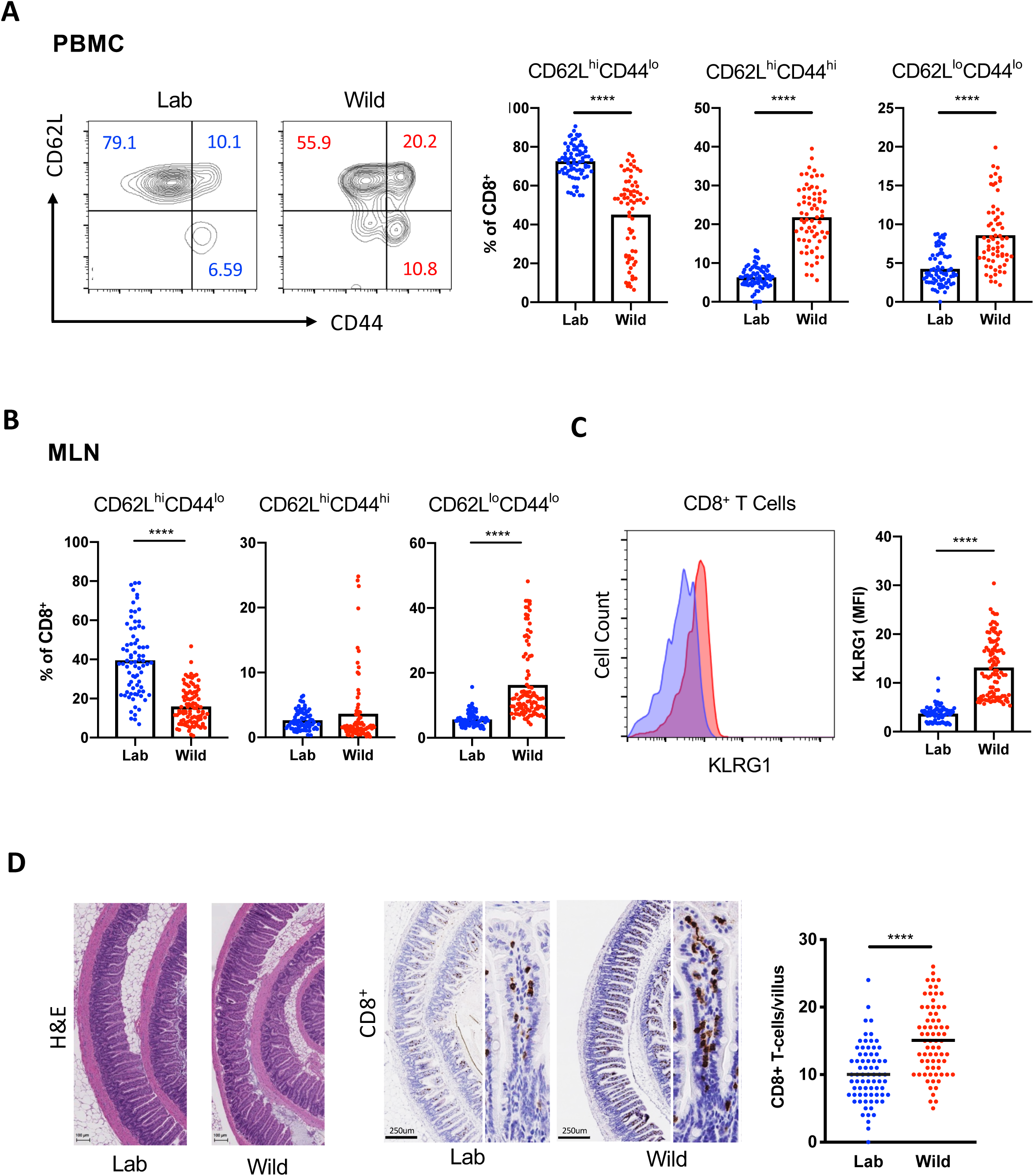
Rewilding alters the proportion of lymphoid cell subsets. (**A**) Representative flow cytometry plots and quantification of CD62L^hi^CD44^lo^, CD62L^hi^CD44^hi^ and CD62L^lo^CD44^hi^ CD8^+^ T cells in the peripheral blood of rewilded mice (Wild) and control mice maintained in the laboratory condition (Lab). Cells were gated on Live^+^CD45^+^Myeloid^-^ CD3^+^CD19^-^CD8^+^. N = 79 lab and 101 rewilded mice (**B**) Quantification of the indicated CD8^+^ T cells in the mesenteric lymph nodes (MLNs). (**C**) Mean fluorescent intensity of MLN CD8^+^ T cells expressing activation marker KLRG1. (**D**) Representative images of small intestinal sections stained with H&E or anti-CD8 antibodies and quantification. N = 50 villi from 5 mice per condition. **** *P* < 0.0001 by two-tailed Student’s *t*-test between groups, (A) to (D).

### Rewilding leads to an increase in granulocytes

We complemented the above flow cytometric and microscopic analyses of T cells by performing RNA-Seq on MLNs from the same lab and rewilded mice described above. 112 genes were differentially regulated by >1.5-fold, the majority of which have known immune-related functions and were upregulated in the rewilded condition (Fig. 2A). Consistent with the aforementioned changes in T cells, pathway analyses indicated that release into an outdoor enclosure increased the expression of genes involved in lymphocyte signaling, differentiation, and migration. Other differentially regulated genes represented pathways associated with polarization of myeloid cell types, such as IL-12 production and iNOS signaling (Fig. 2A). Wild mice have been shown to harbor changes in myeloid sub-populations (Abolins et al., 2017). Therefore, we examined the flow cytometry data for potential changes to non-lymphocyte cell types. We found a striking increase in the side scatter high (SSC^hi^) fraction of PBMCs and cells harboring myeloid markers in rewilded mice compared with lab mice, indicating that release into the outdoor enclosure induced an enrichment in granulocytes (Fig. 2B). Similarly, we observed an increase in both the proportion and total numbers of neutrophils in the MLNs of rewilded mice (Fig. 2C). These findings indicate that rewilding leads to substantial changes in granulocyte numbers and identifies expansion of this leukocyte subset as another key difference in addition to differentiated T cells between the immune system of lab mice and mice exposed to the natural environment.

**Figure 2.**
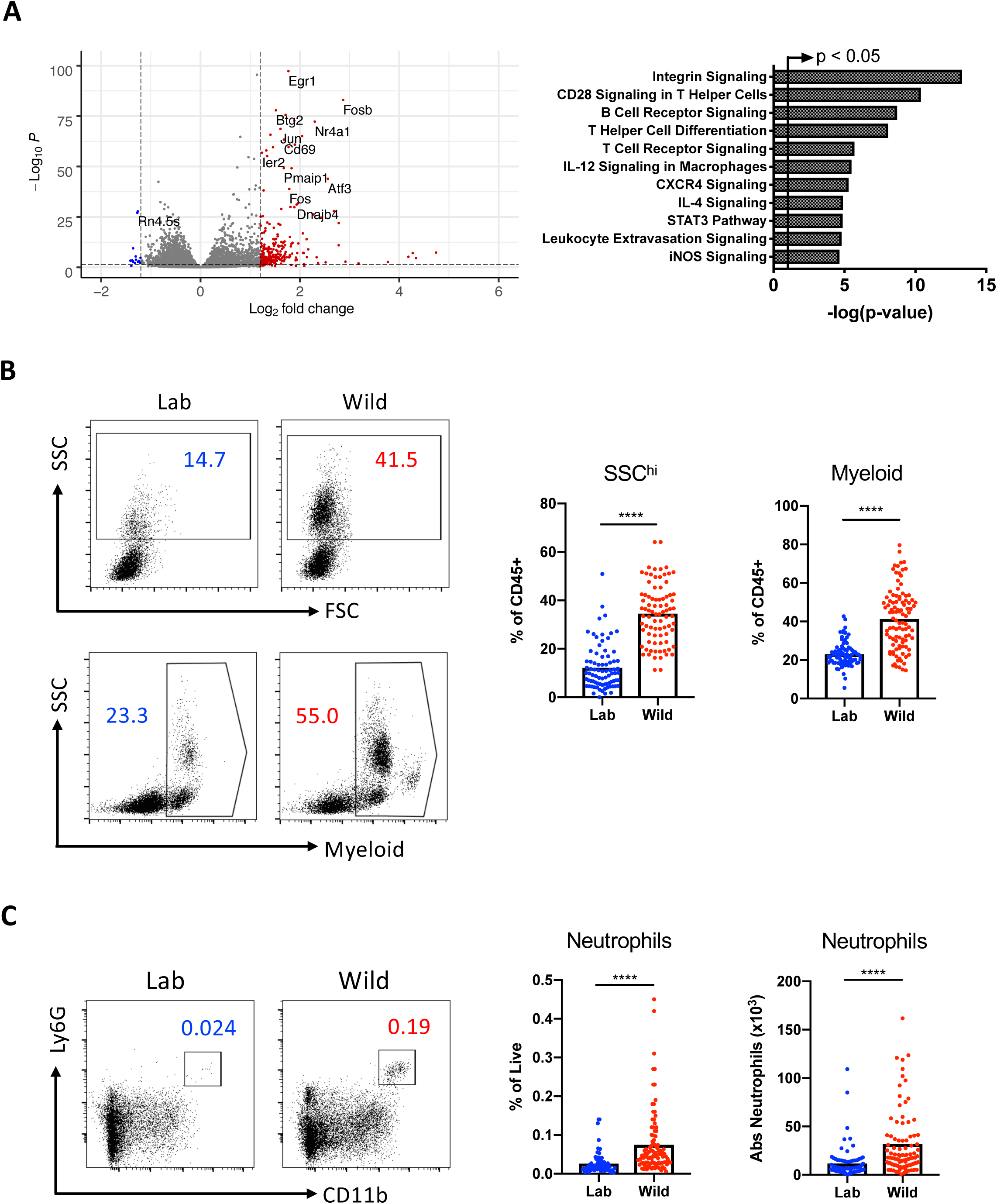
Rewilding leads to an expansion in granulocytes. (**A**) Functional classification of 112 differentially regulated genes (log_2_FC>1.5) by Ingenuity pathway analysis comparing RNA-Seq of MLN cells from lab and rewilded mice. The arrowed line marks where the p-value becomes less than 0.05. N = 40 lab and 46 rewilded mice. (**B**) Representative forward scatter (FSC) and side scatter (SSC) flow cytometry plots and quantification of SSC^hi^ cells (granulocytes) and myeloid cells in the peripheral blood of lab and rewilded mice. All cells were gated on Live^+^CD45^+^ and myeloid cells were identified by cell surface markers CD11b/CD11c/DX5. N = 79 lab and 101 rewilded mice. (**C**) Quantification of the absolute number of MLN cells expressing neutrophil markers (CD11b^+^Ly6G^+^). **** *P* < 0.0001 by two-tailed Student’s *t*-test between groups, (B) to (C).

### Lymph node cells from rewilded mice display enhanced responses to microbial antigens

The ability of immune cells to respond to microbes can be assessed by measuring cytokine production upon stimulation with antigens *ex vivo*. At the time of sacrifice, we distributed single cell suspensions of MLNs from rewilded and lab mice into 96-well plates pre-coated with a panel of UV-killed microbes (Fig. 3A) that are frequently found in the soil, surfaces of the human body, or both: *Bacillus subtilis*, *Pseudomonas aeruginosa*, *Clostridium perfringens*, *Candida albicans*, *Bactroides vulgatus*, and *Staphylococcus aureus*. We also included PBS as a negative control to establish baseline production of soluble factors and αCD3/CD28 beads as a positive control that non-specifically activates T cells. A multiplex bead array was used to quantify 13 cytokines and chemokines in the supernatant following 48-hour incubation.

**Figure 3.**
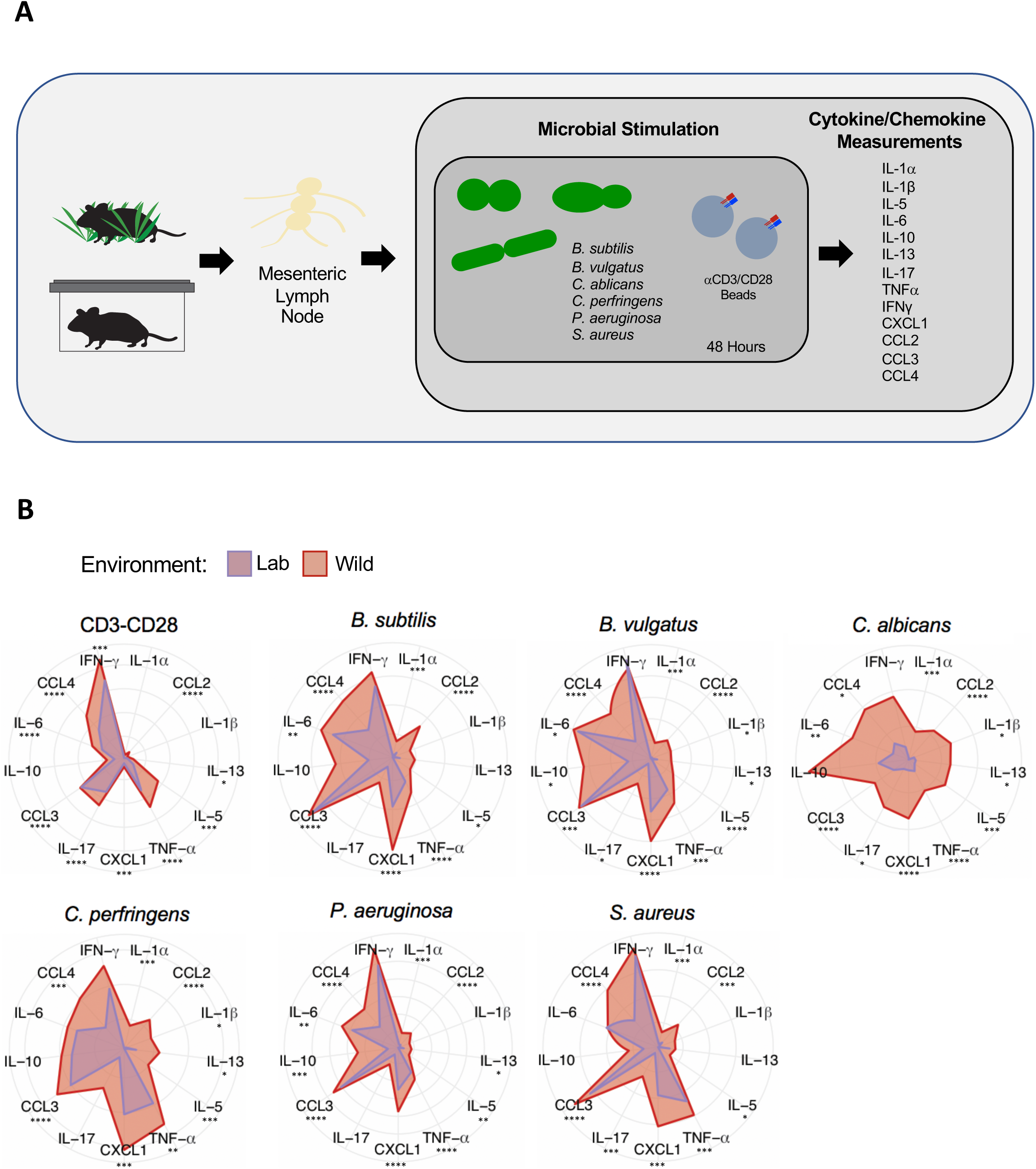
Lymph node cells from rewilded mice display increased cytokine production in response to microbial stimulation. (**A**) Schematic of the experimental design: single cell suspensions of MLNs from lab and rewilded mice were distributed in 96 well plates containing αCD3/CD28 beads or the following microbes: *Bacillus subtilus*, *Bacteroides vulgatus*, *Candida albicans*, *Clostrdium perfringens*, and *Staphylococcus aureus*. The indicated cytokines and chemokines were measured in the supernatant after 48 hours of stimulation. (**B**) Radar plots showing the amount of each indicated cytokine/chemokine produced in response to the microbial stimulants. Data represents Log2 fold change over MLN cells unstimulated PBS controls. N = 40 lab and 47 rewilded mice. * *P* < 0.05, ** *P* < 0.01, *** *P* < 0.001, **** *P* < 0.0001 by two-tailed Student’s *t*-test between groups, (B).

MLN cells derived from rewilded mice responded to microbial or T cell stimulation by secreting generally higher amounts of cytokines and chemokines compared with cells derived from lab mice (Fig. 3B). The exact cytokines that were over-produced were specific to the microbial stimulant, although some general trends include 2–10-fold increases in IL-17, CXCL1, CCL3, and IL-10. *Ex vivo* stimulation with the model fungus *C. albicans*, which frequently colonizes the gastrointestinal tract of humans, led to a particularly striking difference between cells from rewilded versus lab mice. These results suggest that immune cells of rewilded mice are in a hyperactivated state due to exposure to microbes, including fungi.

### Rewilded mice display differential colonization by microbes

Pet store mice and wild mice harbor a multitude of pathogens, some of which are lethal when transmitted to co-housed lab mice (Beura et al., 2016). Rewilded mice are predicted to display reduced exposure to disease-causing pathogens because our enclosure prevents contact with wild rodents. We did not trap a single rodent other than our released lab animals during the entirety of the experiment. To screen for pathogen exposure, we subjected representative mice to a comprehensive serology panel similar to the one that identified the presence of infections in pet store mice (Beura et al., 2016). Rewilded mice were sero-negative for all 24 of the pathogens tested in this panel (Table S1A). Also, an extended PCR-based analysis that includes a skin swab was negative for most agents (see below), and we did not detect Tritrichomonas protozoan species recently shown to mediate expansion of granulocytes in mice (Chudnovskiy et al., 2016) (Table S1B).

Through the above PCR-based assay, several agents were detected in a subset of rewilded mice (15-95%) that may be considered opportunistic pathogens, although these microbes are often described as commensals that are part of the microbiota depending on the context. *Staphylococcus aureus* was detected in the stool of most of the rewilded mice examined. *S. aureus* is a commensal bacterium of the skin in humans and typically not considered a gastrointestinal pathogen, although fecal-oral transmission may be important for spread of antibiotic resistant strains associated with invasive diseases in humans and can be found in the intestines of mice following experimental inoculation (Kernbauer et al., 2014b). Sequencing of the *S. aureus* strain we isolated from a representative rewilded mouse revealed the presence of an intact *β-hemolysin* gene that was not disrupted by prophage φSa3, which encodes immune modulatory proteins that are highly human specific, indicating that rewilded mice are likely colonized by an animal strain rather than a clinical isolate (McCarthy et al., 2012; Price et al., 2012) (Table 2). We found that a few of the mice within our SPF facility harbored *S. aureus*, suggesting that this bacterium was already present in a subset of mice prior to release into the outdoor enclosure and was spread within the enclosure. Thus, the presence of *S. aureus* is not unique to the rewilding conditions. The PCR panel also indicated that a subset of rewilded mice harbored other microbes that are frequently found in SPF facilities, such as *Proteus* and *Klebsiella* species that belong to the Enterobacteriaceae family of pathobionts (Table S1B). We were able to isolate Enterobacteriaceae species including *Citrobacter koseri* from stool of rewilded mice plated on MacConkey agar, which selects for Gram-negative bacilli from the gut.

**Table 2.**
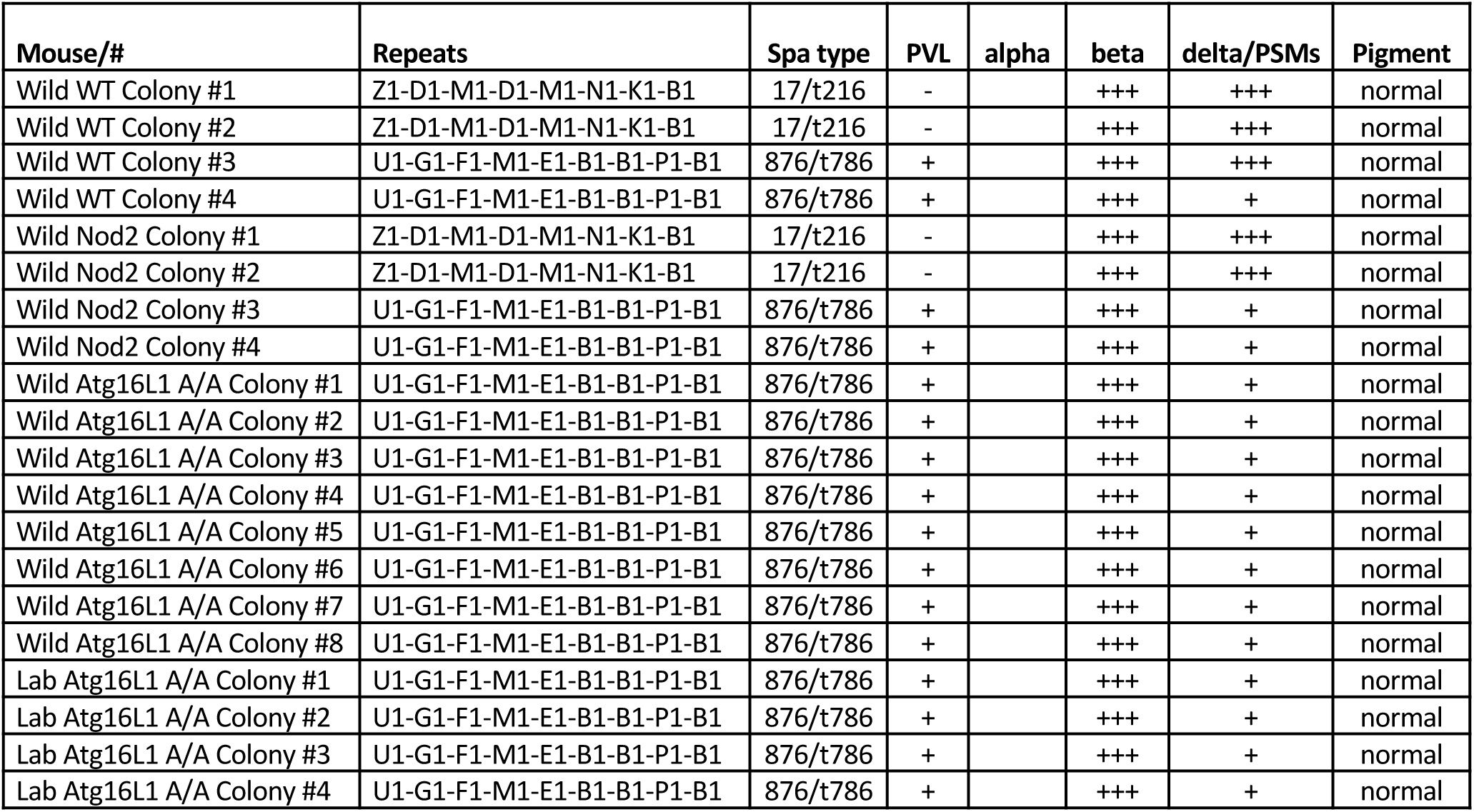
Staphylococcus aureus isolates identified from stool of rewilded mice. Genotyping of S. aureus isolates from representative lab and rewilded mice by DNA sequence analysis of the protein A gene variable repeat region (spa typing) and a variety of additional DNA polymorphisms including the presence of pvl genes.

We inoculated mice with pooled Enterobacteriaceae species isolated from rewilded mice, and although we detected colonization, T cell and granulocyte population were unchanged (fig. S2, A to B). Therefore, *Enterobacteriaceae* species are unlikely to be sufficient to mediate the changes to the immune system observed in rewilded mice.

We next turned our attention to other components of the microbiota as a potential source of immune activation. 16S rDNA sequencing of stool samples indicated that rewilding was associated with enrichment of *Bacteroides* and related taxa, and a decrease in Firmicutes such as *Lactobacillus* and *Faecalibaculum* species (Fig. 4A-D). However, these changes were modest when considering the previously reported large increase in *Bacteroides* abundance in the microbiota of wild mice compared with SPF mice (Rosshart et al., 2017), likely reflecting the fact that in our system, adult mice are released into the wild environment with an intact microbiota. Although we did not detect a change in the number of taxa represented in rewilded mice compared with lab mice (Fig. 4B), it is possible that the coding capacity of the microbiota is altered following release into the outdoor enclosure. Therefore, we performed shotgun sequencing on a subset of WT mice. Principle coordinate analysis (PCA) indicated that the community of microbes segregate by environment (lab versus wild) (Fig. 4E). 184 gene families were identified as differentially regulated, many of which were attributed to the *Parabacteroides* genus and are associated with metabolic processes (fig. S3A). Together with the 16S sequencing results, these findings indicate that the composition and gene content of the bacterial microbiota is different in lab and rewilded mice.

**Figure 4.**
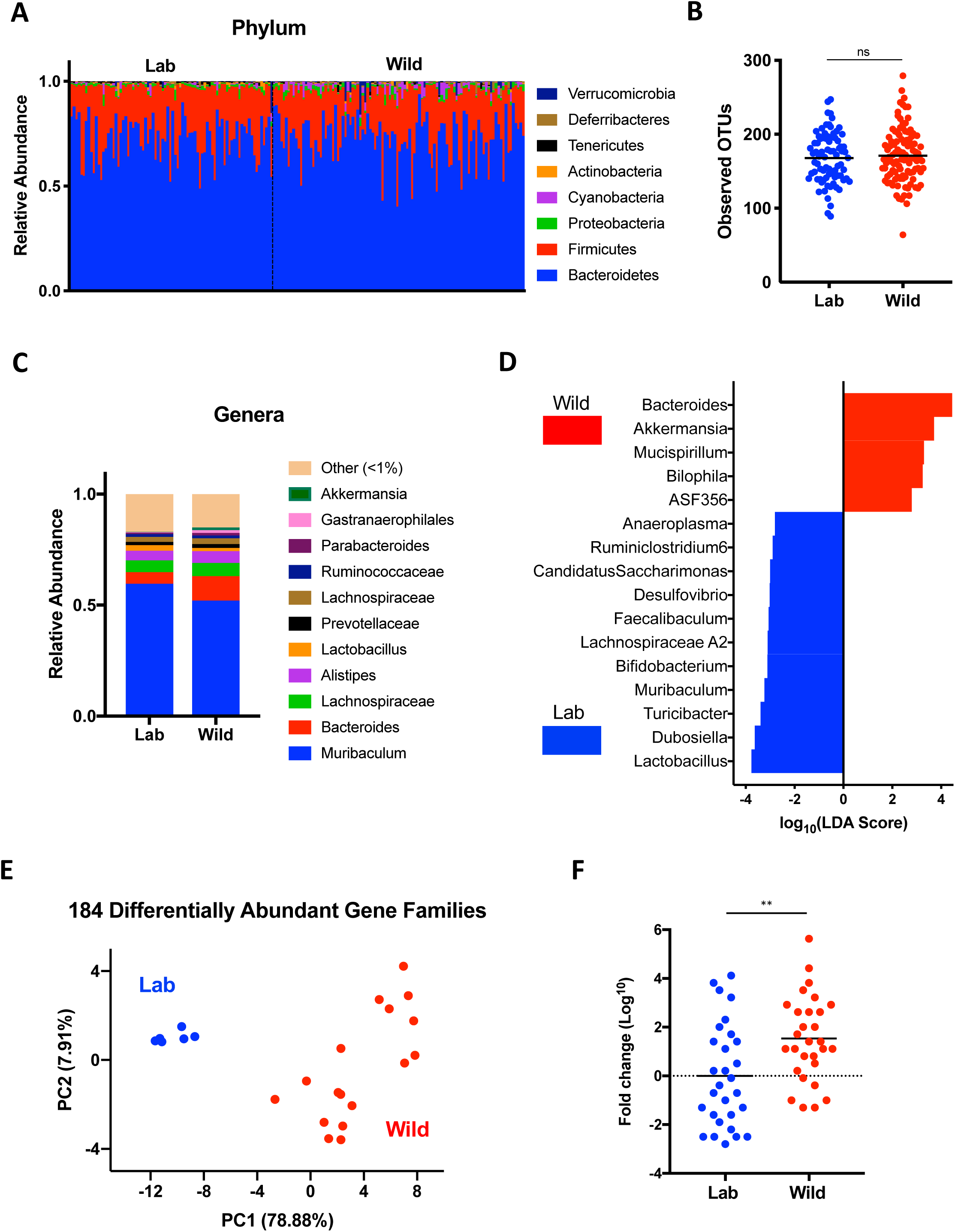
Altered immunity in rewilded mice is asso ciated with microbial exposure in the natural environment. (**A**) Relative abundance of phylum level taxa that constitute the fecal microbial community of lab (Lab) and rewilded (Wild) mice as determined by 16S sequencing. N = 79 lab and 102 rewilded mice. **(B)** Alpha diversity quantification through number of operational taxonomic units (OTUs). (**C**) Stacked bar plot of mean relative abundances of genus level taxa. **(D)** Bacterial taxa from (C) significantly enriched in lab versus wild conditions as determined by linear discriminant analysis effect size (LEfSe) analysis using a LDA threshold score of 2.5. (**E**) Principle coordinate analysis of differential microbial gene families abundance between lab and wild mice determined by shotgun sequencing. N = 6 lab and 17 rewilded mice(**F**) Quantification of relative fungal burden in stool of lab and rewilded mice as determined by qPCR of the internal transcribed space (ITS) region normalized to the average of lab mice. N = 45 lab and 50 rewilded mice. **** P <0.01 by two-tailed Student’s *t*-test between groups, (F).

Our experiments in which we quantified cytokine production by antigen-stimulated MLN cells suggest that rewilding is associated with reactivity to fungi. We mined the shotgun sequencing reads for fungal genomes and found that samples from rewilded mice included sequences corresponding to fungal taxa such as *Candida* (fig. S3B). Quantification of fungal burden by qPCR of the conserved internal transcribed spacer (ITS) region of ribosomal RNA on the same samples as above indicated that rewilded mice harbor a significant increase in intestinal colonization by fungi compared with lab mice (Fig. 4F). These findings raise the possibility that acquisition of fungi contribute to changes in immunity. Below, we analyze the effect of *Atg16l1* and *Nod2* mutations on immune parameters, and subsequently examine the effect of introducing fungi into lab mice.

### *Atg16L1* and *Nod2* mutations are associated with signs of disease but do not affect immune cell populations

*ATG16L1* and *NOD2* are among the most highly investigated susceptibility genes for IBD, an inflammatory disorder of the gut that is associated with alterations in the microbiota (Wlodarska et al., 2015). Inhibition of ATG16L1, which mediates the cellular degradative process of autophagy, leads to excess inflammasome activity and epithelial defects in models of intestinal inflammation (Cadwell et al., 2008; Cadwell et al., 2010; Lassen et al., 2014; Murthy et al., 2014; Saitoh et al., 2008). NOD2 is a cytosolic bacterial sensor that is important for host defense, as demonstrated by the heightened sensitivity of *Nod2^-/-^* mice to inflammation and mortality during oral infection by the extracellular Gram-negative pathogen *Citrobacter rodentium* (Kim et al., 2011). Although NOD2 can function upstream of ATG16L1 to promote immunity against intracellular bacteria or antigen presentation (Chu et al., 2016; Cooney et al., 2010; Travassos et al., 2010), *Atg16l1* mutant mice display enhanced immunity towards *C. rodentium* (Marchiando et al., 2013; Martin et al., 2018), indicating that mutations in these genes can lead to different outcomes when responding to microbial colonization of the gut. Therefore, we included both mice harboring the IBD variant of *Atg16l1* (T316A) and *Nod2^-/-^* mice in the cohort of animals released into the enclosure.

Only a subset of mice in the enclosure displayed any sign of disease. Rewilded *Atg16l1^T316A^* homozygous (5 out of 24) and heterozygous mice (1 out of 27) presented with diarrhea, and none of the other genotypes of rewilded mice or lab mice showed watery stool. Interestingly, all the *Atg16l1* mutant mice with diarrhea were collected during the second week of the capture (Fig. 5A). Although diarrhea was not observed in *Nod2^-/-^* mice at any point during the experiment, many of them failed to gain weight following release into the enclosure (Fig. 5B). Despite these observations, we did not detect intestinal inflammation by histopathology in small intestinal or colonic tissue sections under any condition (data not shown).

**Figure 5.**
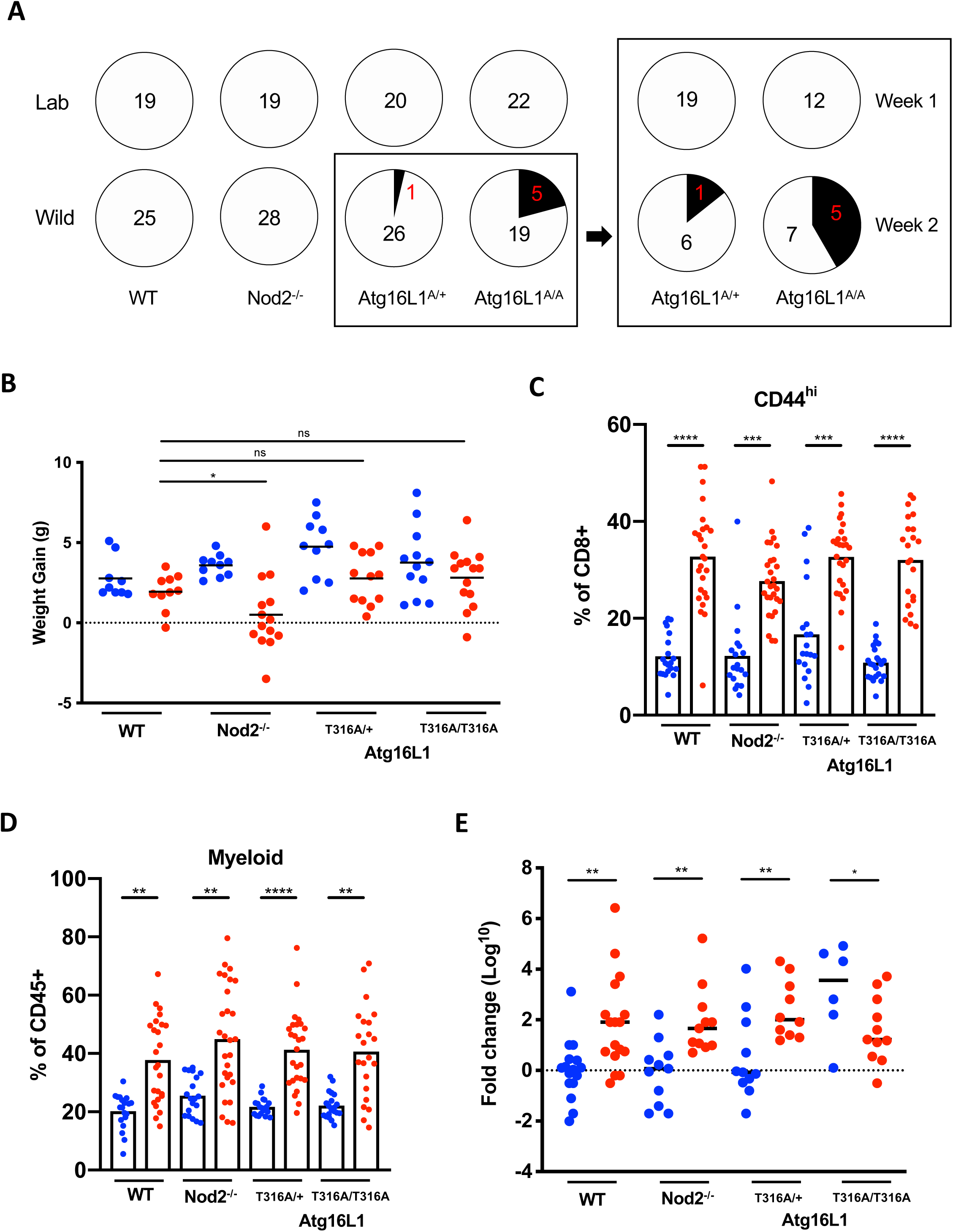
Atg16L1 and Nod2 variants are associated with disease pathology but do not affect immune cell populations. **(A)** Number of mice presenting diarrhea at time of sacrifice for lab and rewilded wildtype (WT), *Atg16l1*, and *Nod2* mutant mice. Right panel displays number of *Atg16l1^T300A^* het (A/+) and homozygous (A/A) mice with diarrhea when sacrificed during the first week or second week of trapping. N > 11 mice per group **(B)** Weight gain of the WT, *Atg16l1*, and *Nod2* mutant mice over a period of 7 weeks in Lab (blue) and Wild (red) conditions. N > 20 mice per group (**C**) Quantification of CD44^+^CD8^+^ T cells in the peripheral blood of WT, *Atg16l1*, and *Nod2* mutant mice in Lab (blue) and Wild (red) conditions. **(D)** Quantification of myeloid cells as identified by cell surface markers CD11b/CD11c/DX5. **(E)** Quantification of relative fungal burden in stool of WT, *Atg16l1*, and *Nod2* mutant mice in Lab (blue) and Wild (red) conditions as determined by ITS qPCR. N > 6 mice per group. * *P* < 0.05, ** *P* < 0.01, *** *P* < 0.001, **** *P* < 0.0001 by two-tailed Student’s *t*-test between groups, (B) to (E).

Next, we examined whether mutation in these IBD genes affect the expansion of differentiated T cells and myeloid cells, the focus of this study. We found that these leukocyte populations in the blood displayed similar increases in rewilded *Atg16l1* and *Nod2* mutant mice as their wild-type counterparts (Fig. 5C-D). We also examined whether *Atg16l1* and *Nod2* mutant mice differ in their microbiota composition. The results from breaking down the 16S analyses by genotype were similar to those from analyses of the pooled data in Figure 3. Generally, genera that were enriched or reduced in rewilded WT mice were also enriched or reduced in rewilded *Atg16l1* and *Nod2* mutant mice, and the total number of observed taxa were similar across all conditions (fig. S4A-B).

The more striking effect of genotype was on fungal colonization. We found that lab *Atg16l1* mutant mice displayed expansion of fungi to levels similar to rewilded mice (Fig. 5E). This expansion in the lab environment potentially reflects the role of ATG16L1 in the antifungal pathway of LC3-associated phagocytosis (Akoumianaki et al., 2016; Martinez et al., 2015; Oikonomou et al., 2016). Separating mice by genotype revealed that all of the rewilded mice regardless of genotype had similar levels of fungal colonization. Rewilded *Nod2^-/-^* mice had a similar expansion of fungi as rewilded WT mice. Rewilded WT mice displayed a 150-fold increase in fungal load compared with lab WT mice (Fig. 5E). Given the striking difference in the fungal burden between lab and rewilded WT mice, we focused the remainder of this study on examining the identity of these fungi and their contribution to changes in leukocyte populations.

### Microbial colonization induces granulocytes expansion

The effects of rewilding could be a response to a number of possible environmental variables, including diversification of diet and activity levels outdoors (e.g., (Budischak et al., 2018)). To determine the extent to which the enhanced activation and maturation of the immune system was due to changes in microbial exposure, germ-free mice were reconstituted with microbiota through inoculation with cecal contents from rewilded and lab mice, and then blood and MLNs harvested from their offspring were analyzed by flow cytometry. We chose 3 representative lab and rewilded WT mice (defined as within a standard deviation of the average number of blood granulocytes in the respective conditions) as donors of cecal contents. Mice harboring the microbiota of rewilded mice displayed an increase in granulocytes including neutrophils (Fig. 6, A and B). However, this increase was modest compared with the rewilded mice that served as donors. 16S rDNA sequencing showed that the bacterial community structures were preserved in the offspring of the germ-free mice reconstituted with cecal contents (fig. S5A). In contrast, quantification of total fungal burden indicated that the difference in fungal load within the microbiota of lab versus rewilded mice was not maintained in the progeny of reconstituted mice (Fig. 6C). Therefore, it is possible that sustained exposure to a high fungal burden is necessary to recreate the full effect of rewilding.

**Figure 6.**
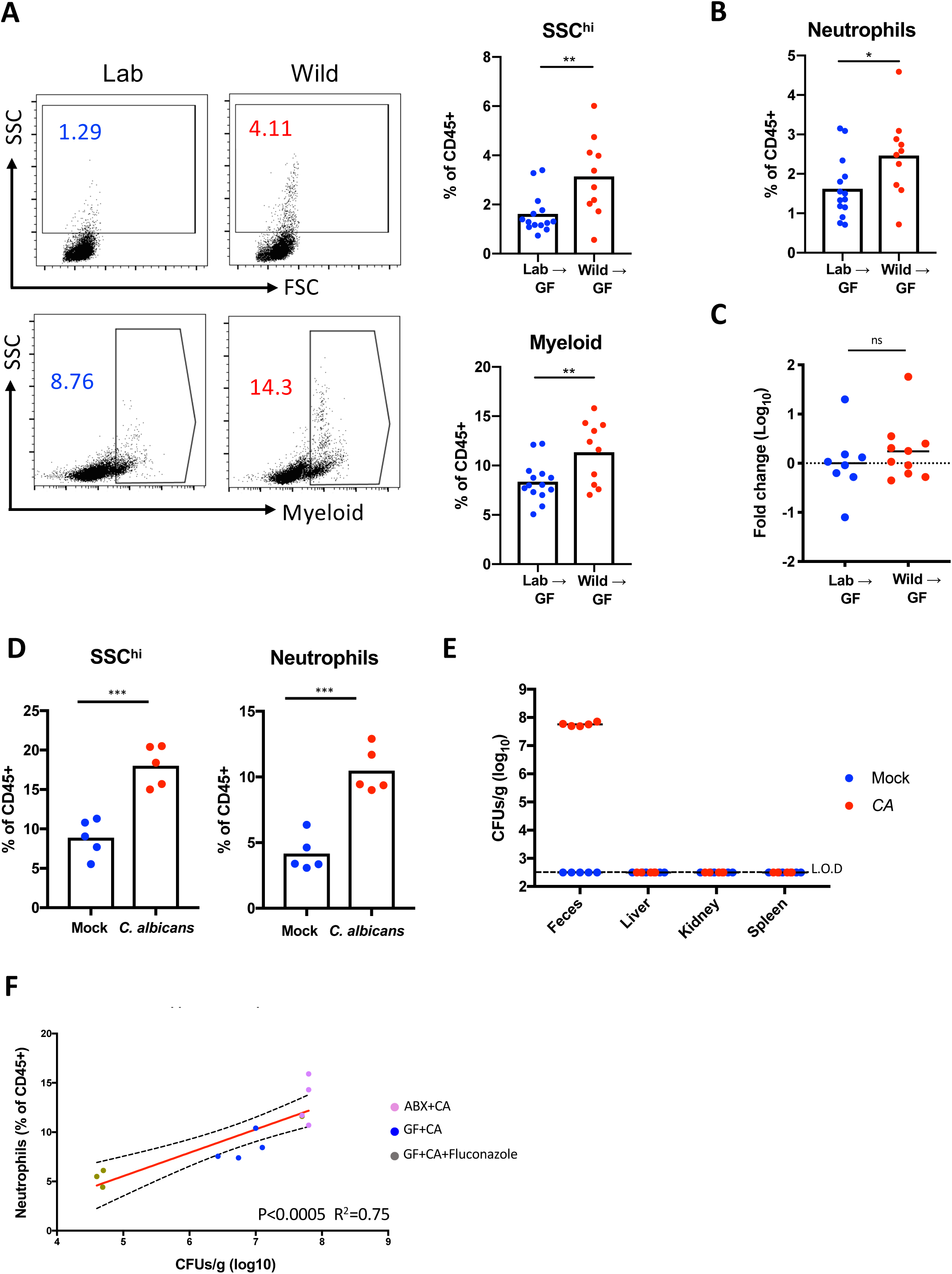
Altering the microbiota of lab mice recreates expansion of granulocytes observed in rewilded mice. (**A**) Representative flow cytometry plots and quantification of SSC^hi^ cells and myeloid cells (CD11b/CD11c/DX5) gated on Live^+^CD45^+^ in the peripheral blood of the F1 progeny from germ-free (GF) mice reconstituted with cecal contents from lab (Lab) and rewilded (Wild) mice. N>10 recipient mice per condition were reconstituted with 3 donor lab and rewilded mice each. Data represents 2 independent repeats. (**B**) Quantification of the proportion of neutrophils (CD11b^+^Ly6G^+^) in the peripheral blood from the mice in (A). (**C**) Quantification of the relative fungal burden in stool of mice from (A) as determined by ITS qPCR. N = 8 and 10 mice reconstituted with lab and rewilded cecal contents from (A), respectively. (**D**) Quantification of SSC^hi^ cells and neutrophils in the peripheral blood from antibiotic-treated conventional mice 4 weeks post-inoculation with PBS or *C. albicans*. N = 5 mice per group, 2 independent repeats. (**E**) Colony forming units (CFUs) of fungi in feces and indicated organs for mice from (D). Dotted line denotes limit of detection (L.O.D.). (**F**) Linear regression analysis comparing frequency of neutrophils in the blood and fungal CFUs in feces from antibiotics treated mice inoculated with *C. albicans*(ABX+CA) from (D), germ-free mice mono-associated with *C. albicans* (GF+CA), and germ-free mice mono-associated with *C. albicans* and treated with fluconazole (GF+CA+Fluconazole). N = 5 ABX+CA, 4 GF+CA, and 3 GF+CA+Fluconazole. * P <0.05, ** P <0.01, *** *P* < 0.001 by two-tailed Student’s *t*-test between groups, (A) to (D).

Intestinal colonization by commensal fungi (the gut mycobiota) can contribute to mammalian immunity and represents an understudied component of the microbiota (Chiaro et al., 2017; Iliev et al., 2012; Jiang et al., 2017; Leonardi et al., 2018; Li et al., 2018; Limon et al., 2019; Tso et al., 2018; Wheeler et al., 2016; Zhang et al., 2016). To circumvent the technical difficulties with establishing prolonged fungal colonization in lab mice, we used an established model in which *C. albicans* is introduced into antibiotics-treated mice (fig. S5B). *C. albicans* was an attractive model to initially test our hypothesis because we demonstrated that MLNs from rewilded mice are hyper-reactive to this fungus (Fig. 3). We found that antibiotics-treated mice inoculated with *C. albicans* displayed a substantial increase in granulocytes and neutrophils compared with similarly treated mice receiving a control inoculum (Fig. 5D). In addition, increases in differentiated CD8^+^ T cell populations were observed in *C. albicans* colonized mice (fig. S5C). These immune cell composition changes were not due to systemic infection as *C. albicans* colonization was restricted to the gut in this model (Fig. 5E). Inoculation of germ-free mice with *C. albicans* also reproduced the increase in granulocytes demonstrating that fungal colonization is sufficient, though this approach led to a lower amount of *C. albicans* colonization, and a corresponding less dramatic change in granulocyte numbers (Fig. 5F and fig. S5D). To further demonstrate that fungal burden directly influences granulocyte numbers, we treated *C. albicans* mono-associated mice with the antifungal drug fluconazole. Our results show that pharmacologically decreasing fungal burden rapidly leads to a further decrease in granulocytes (Fig. 5F and fig. S5E), indicating that the degree of intestinal colonization by fungi impacts the proportion of immune cells in the periphery.

### Fungi isolated from rewilded mice induce granulocyte expansion in lab mice

**(A)** *C. albicans* represents a useful model and is relevant to human disease, but it is unclear whether it is representative of fungi acquired in the outdoor enclosure. Having determined that exposure to the natural environment leads to a large increase in fungal burden, we wished to compare the precise composition of the fungal microbiota of rewilded and lab mice. Fungal sequences represent a minor proportion of shotgun sequencing reads due to the dominant presence of bacteria. Therefore, we performed ITS sequencing of stool to increase our resolution of fungal taxa. We found that the fungal microbiota of rewilded and lab mice segregated by PCA (Fig. 7A and fig. S6A). In contrast to the bacterial microbiota, we found that rewilding was associated with an increase in α-diversity of fungi detected (Fig. 7B). We observed increases in several fungal taxa, most notably an enrichment in *Aspergillus* species (Fig. 7C-D). Therefore, exposure to the natural environment leads to the acquisition of an altered and expanded fungal microbiota.

**Figure 7.**
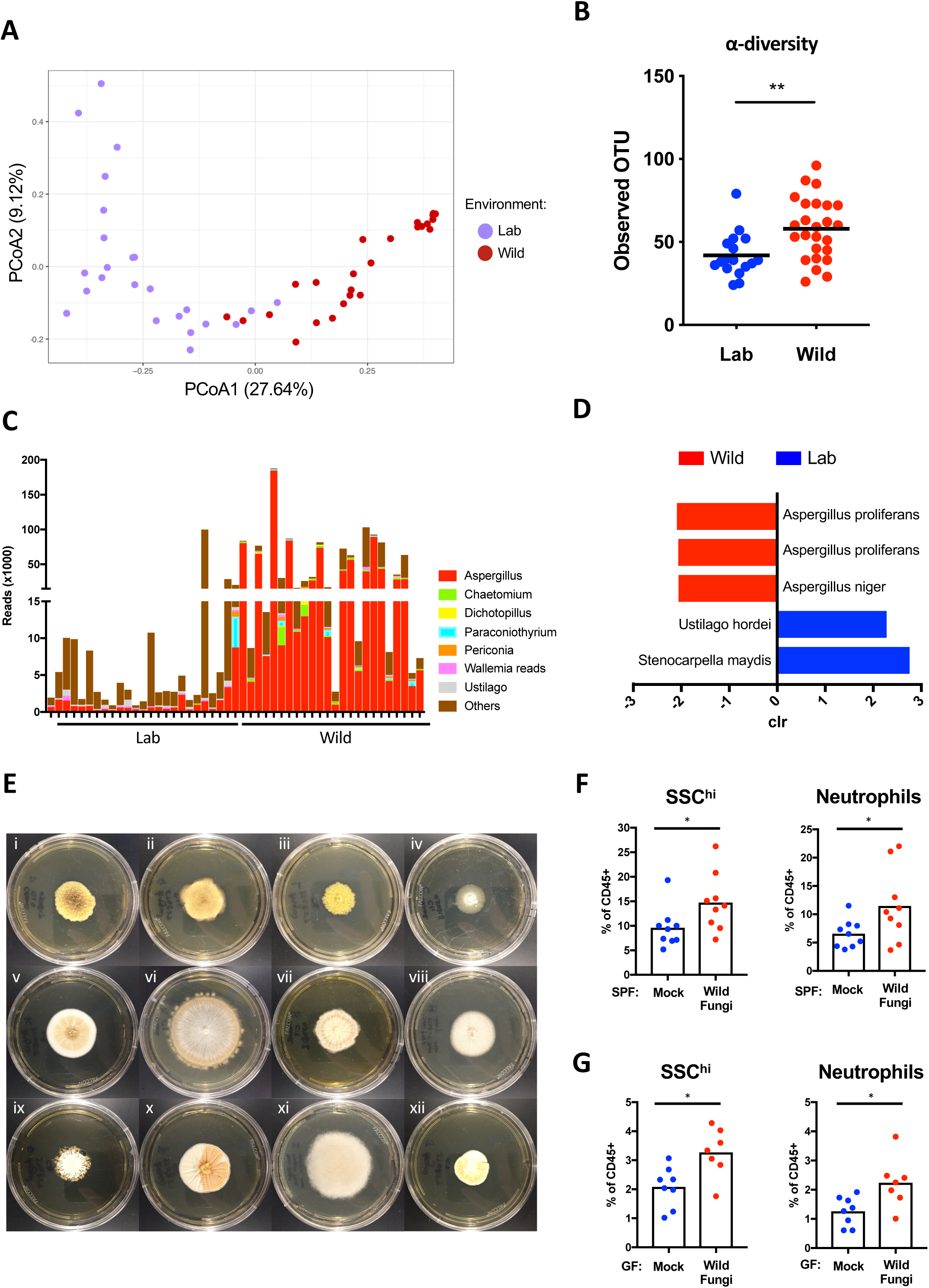
Rewilding leads to colonization by immunomodulatory fungi. (**A**) Beta diversity (Bray Curtis) plots of fungal taxa detected by ITS sequencing of stool from lab (Lab) mice (purple dots) and rewilded (Wild) mice (red dots). N = 24 lab and 25 rewilded mice. (**B**) Alpha diversity quantification through number of operational taxonomic units (OTUs) from (A). (**C**) Number of ITS sequencing reads corresponding to the indicated genus from (A). (**D**) Fungal species from (A) significantly enriched in lab versus wild conditions by analysis of composition of microbiomes (ANCOM). (**E**) Representative images of colonies growing on Sabouraud dextrose agar plates following plating of cecal suspensions harvested from rewilded mice corresponding to (i) *Aspergillus proliferans*, (ii) *Aspergillus proliferans*, (iii) *Aspergillus proliferans*, (iv) *Aspergillus candidus*, (v) *Aspergillus sp.*, (vi) *Chaetomium globosum*, (vii) *Dichotomopilus indicus*, (viii) *Aspergillus niger*, (ix) *Malbranchea flavorosea*, (x) *Talaromyces tratensis*, (xi) *Syncephalastrum racemosum*, (xii) *Talaromyces tratensis*. (**F**) Frequency of SSC^hi^cells and neutrophils in peripheral blood from conventional mice repetitively gavaged with a consortium of 7 wild fungi isolates from plates (i)–(vii) in (E) or PBS for 2 weeks. N = 9 mice per group, 2 independent repeats. **(G)** Frequency of SSC^hi^ cells and neutrophils in the peripheral blood from germ-free mice 2 weeks post-inoculation with a consortium of 7 wild fungi. N > 7 mice per group, 2 independent repeats. * *P* <0.05, ** *P* <0.01, **** *P* < 0.0001 by two-tailed Student’s *t*-test between groups, (B), (F), and (G).

Finally, we examined whether fungi acquired in the outdoor enclosure are capable of driving changes to granulocyte numbers that we observed in rewilded mice. We were able to isolate many of the fungi corresponding to the taxa identified in the ITS sequencing reads when we plated stool from rewilded mice on Sabouraud dextrose agar (Fig. 7E), but were unable to grow any fungi from lab mice. We gavaged SPF mice every other day repeatedly for 2 weeks (fig. S6B) with a consortium of representative “wild” fungi consisting of *Aspergillus candidus*, *Aspergillus proliferans*, *Chaetomium globosum* and *Dichotomopilus Indicus* (corresponding to plates (i)–(vii) depicted in Fig. 7E). We found that these wild fungi were able to induce the increase in peripheral granulocytes and neutrophils (Fig. 7F). To test whether these fungi were sufficient, we inoculated germ-free mice with the same consortium and found a similar increase in granulocytes and neutrophils (Fig. 7G). Together, these experiments show that fungi in the natural environment can contribute to the state of the immune system.

## DISCUSSION

Recent studies have shown that infection with disease-causing pathogens that are excluded from SPF facilities lead to large-scale changes in immunity that can alter subsequent immune responses (Abolins et al., 2017; Beura et al., 2016; Reese et al., 2016; Rosshart et al., 2019). Exposure to these pathogens have been suggested to ‘correct’ the immune system by increasing the number of mature immune cells to more closely resemble the state of the adult human immune system (Masopust et al., 2017; Tao and Reese, 2017). These fundamentally important findings suggest that childhood infections that occur in most individuals contribute to the education and function of our immune system. However, the extent to which commensal or environmentally-acquired agents are capable of driving these changes in immunity was unclear.

We found that releasing laboratory mice into a natural environment, even for a limited amount of time, can reproduce the enhanced differentiation of memory T cells previously attributed to infection by life-threatening pathogens (Beura et al., 2016). ∼25% of lab mice co-housed with pet store mice did not survive the procedure (Beura et al., 2016), whereas 90% of mice released into our outdoor enclosure were recovered at the end of the 6-7 week period, with several of the stragglers successfully trapped thereafter (all together, >95% recovery). Comprehensive serology indicated that rewilded mice were not infected by the pathogens detected in pet store mice, although we acknowledge the possibility that agents not covered by the assay may be present, and that our rewilded mice were likely exposed to pathobionts that can cause disease in immunocompromised mice. Consistent with the minimal infection by disease-causing pathogens, visual inspection and handling of rewilded WT mice indicated that they were healthy and their movements were sharper and their muscles stronger than lab control mice. Thus, our outdoor enclosure system may be useful for identifying immunomodulatory microbes that are difficult to isolate from wild or pet shop mice that have a more complex infectious history. We suggest that controlled release into nature can complement other approaches that expose mice to the natural environment, and that each method has its strengths and weaknesses.

The most striking effect of rewilding that we identified was the expansion of granulocytes in the blood including neutrophils. This finding is notable because granulocytes are abundant in human blood and scarce in laboratory mice. Hence, we were eager to identify the environmental variables that drive granulocyte expansion. The microbiota of rewilded mice was characterized by a significant increase in total fungi that included acquisition of *Aspergillus* species. Transfer of microbiota from rewilded mice into germ-free mice was able to reproduce the increase in granulocytes. Still, the proportion of blood granulocytes in reconstituted germ-free mice was lower than that of rewilded mice, potentially due to the poor engraftment of fungi. Therefore, we examined the contribution of fungi directly and found that granulocytes expansion could be recreated through inoculation of lab mice with a model human commensal fungus, *C. albicans*, or a cocktail of fungi isolated from rewilded mice. Our observation that the degree of fungal burden recovered in stool correlates with neutrophil numbers provides additional evidence that fungi contribute to the proportion of granulocytes in the blood. Intestinal colonization by *C. albicans* occurs in the majority of adult humans and is associated with enhanced Th17 CD4^+^ T cell differentiation, and IL-17 produced by these T cells can explain the increase in neutrophils (Bacher et al., 2019; Shao et al., 2019). It is possible that the normalization of the immune system that occurs upon rewilding, at least with respect to granulocytes, mimics intestinal colonization by *C. albicans* in humans. We suggest that restoring the fungal microbiota of lab mice can make the murine model a better match to human physiology.

We also had an opportunity to determine the contribution of two IBD susceptibility genes. Neither *Atg16l1* nor *Nod2* mutation had a noticeable effect on the leukocyte populations we measured. Interestingly, *Atg16l1* mutant mice that served as the lab controls displayed a large increase in fungal burden. Although unclear if related, a subset of the rewilded *Atg16l1* mutant mice displayed diarrhea. We previously demonstrated that *Atg16l1* and *Nod2* mutant mice develop small intestinal defects in a manner dependent on specific commensal-like agents, and that close relatives of these agents do not trigger any abnormalities (Cadwell et al., 2010; Kernbauer et al., 2014a; Matsuzawa-Ishimoto et al., 2017; Ramanan et al., 2016; Ramanan et al., 2014). The observation that most of the *Atg16l1* and *Nod2* mutant mice did not display obvious signs of intestinal inflammation may reflect this specificity of gene-microbe interactions in IBD. As we only included two genotypes and on one background (C57BL/6J) in this study, a more comprehensive examination of genetic susceptibility to a change in environment is warranted.

In conclusion, our results indicate that fungi are among the different infectious entities that mammals encounter in the natural environment and represent one of the missing or altered components of the microbiota in lab mice kept under SPF conditions. Microbes, vegetation, and climate vary across the globe, and it is likely that exposure to nature has a different consequence depending on the location. Nevertheless, the recent study characterizing wildings noted that wild mice when compared with lab mice indeed display increased colonization by Ascomycota, the phylum that includes the *Aspergillus* species we found in our rewilded mice (Rosshart et al., 2019). Therefore, exposure to fungi may not be unique to our experimental system, and it remains possible that this fungal exposure drives the altered immunity observed in other studies examining wild or pet shop mice. When taken together, the findings we present here expand our understanding of the immunomodulatory role of intestinal fungi and indicate that a diverse fungal microbiota, likely together with the bacterial microbiota, participates in the differentiation of the immune system in a free-living mammal.

## Supporting information

STAR METHODS

## Acknowledgements

We wish to thank William Craigens, Daniel Navarrete Prado, Allison Lee, and Veena Chittamuri for assistance with trapping and husbandry in the field, the PU Lab Animal Resources staff for logistical support. We wish to thank the NYU School of Medicine Flow Cytometry and Cell Sorting, Microscopy, Genome Technology, and Histology Cores for use of their instruments and technical assistance (supported in part by National Institute of Health (NIH) grant P31CA016087, S10OD01058, and and S10OD018338). We also wish to thank Margie Alva, Juan Carrasquillo, and Beatriz Delgado for technical assistance with gnotobiotics.

## Author contributions

Design of experiments, data analysis, data discussion, and interpretation: F.Y., Y.H.C., J.D.L., J.C.D., P.L., A.L.G., and K.C; primary responsibility for execution of experiments: F.Y., Y.H.C., J.D.L., J.M.L., C.M., A.C., Z.S., and C.D.D.; MLN cell RNA, 16S and ITS sequencing analysis: J.C.D., J.L.R., K.V.R., Z.S., F.Y., and Y.H.C. All authors discussed data and commented on the manuscript.

## Data and materials availability

Raw sequence data from 16*S*, ITS, and RNA sequencing experiments are deposited in the NCBI Sequence Read Archive under BioProject accession number PRJNA559026 and gene expression omnibus (GEO) accession number GSE135472.

---

**Supplemental Figure 1.**
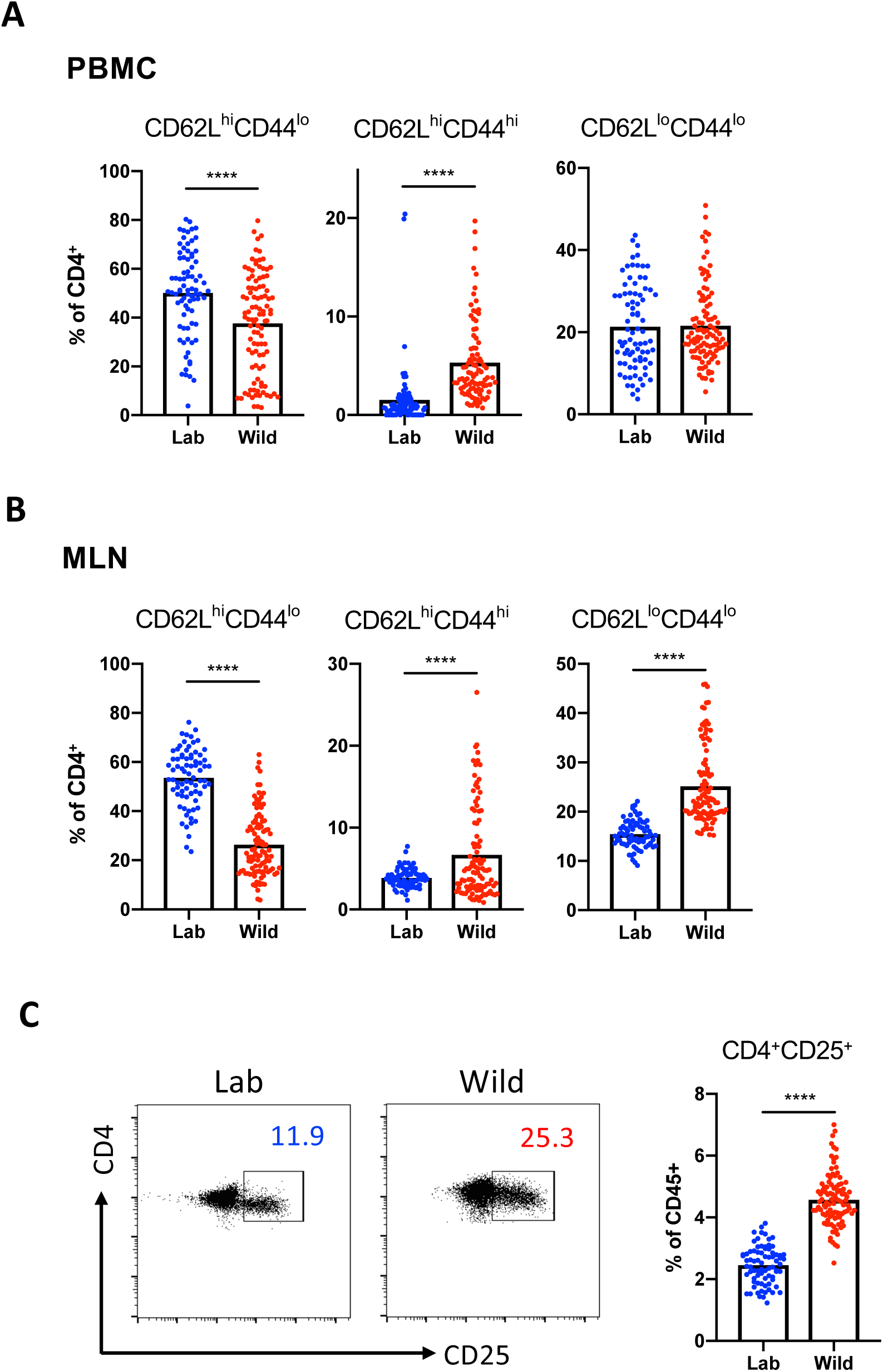
Rewilding increases the proportion of differentiated CD4^+^ T cell populations. Additional quantification of T cell subsets from rewilded (Wild) mice and control mice maintained in the laboratory condition (Lab) described in Figure 1. (**A-B**) Quantification of CD62L^hi^CD44^lo^, CD62L^hi^CD44^hi^, and CD62L^lo^CD44^hi^ CD4^+^ T cells in the (A) peripheral blood and (B) MLNs. (**C**) Representative flow cytometry plots and quantification of mesenteric lymph node (MLN) cells expressing CD4^+^CD25^+^. N = 79 lab and 101 wild mice (Blood); N = 77 lab and 101 rewilded mice (MLN). **** *P* < 0.0001 by two-tailed Student’s *t*-test between groups, (A) to (C).

**Supplemental Figure 2.**
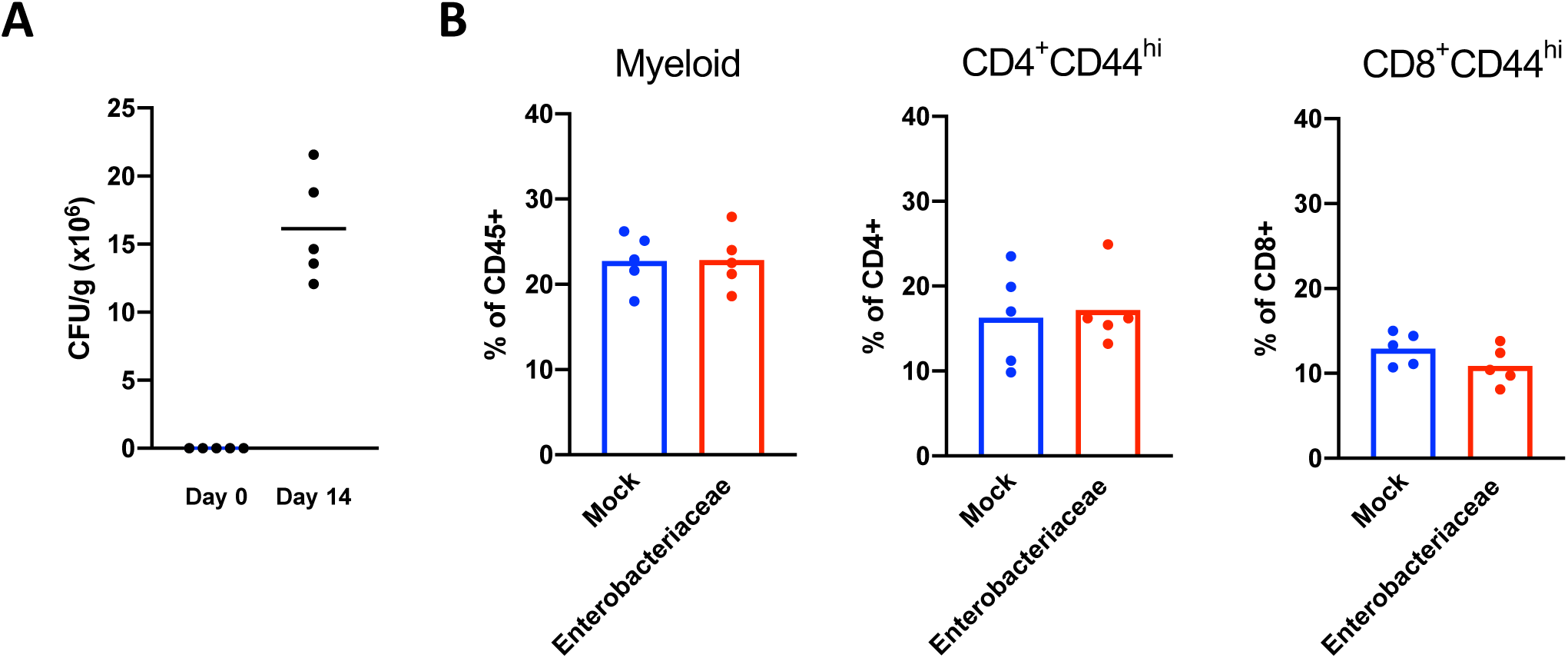
Enterobacteriacae isolated from rewilded mice does not alter immune cell populations. (**A**) Enterobacteriacae burden in the stool of SPF mice before and after colonization (**B**) Quantification of CD11b/CD11c/DX5 (Myeloid), CD4^+^CD44^+^ T cells, and CD8^+^CD44^+^ T cells in the peripheral blood of enterobacteriacae colonized and control mice.

**Supplemental Figure 3.**
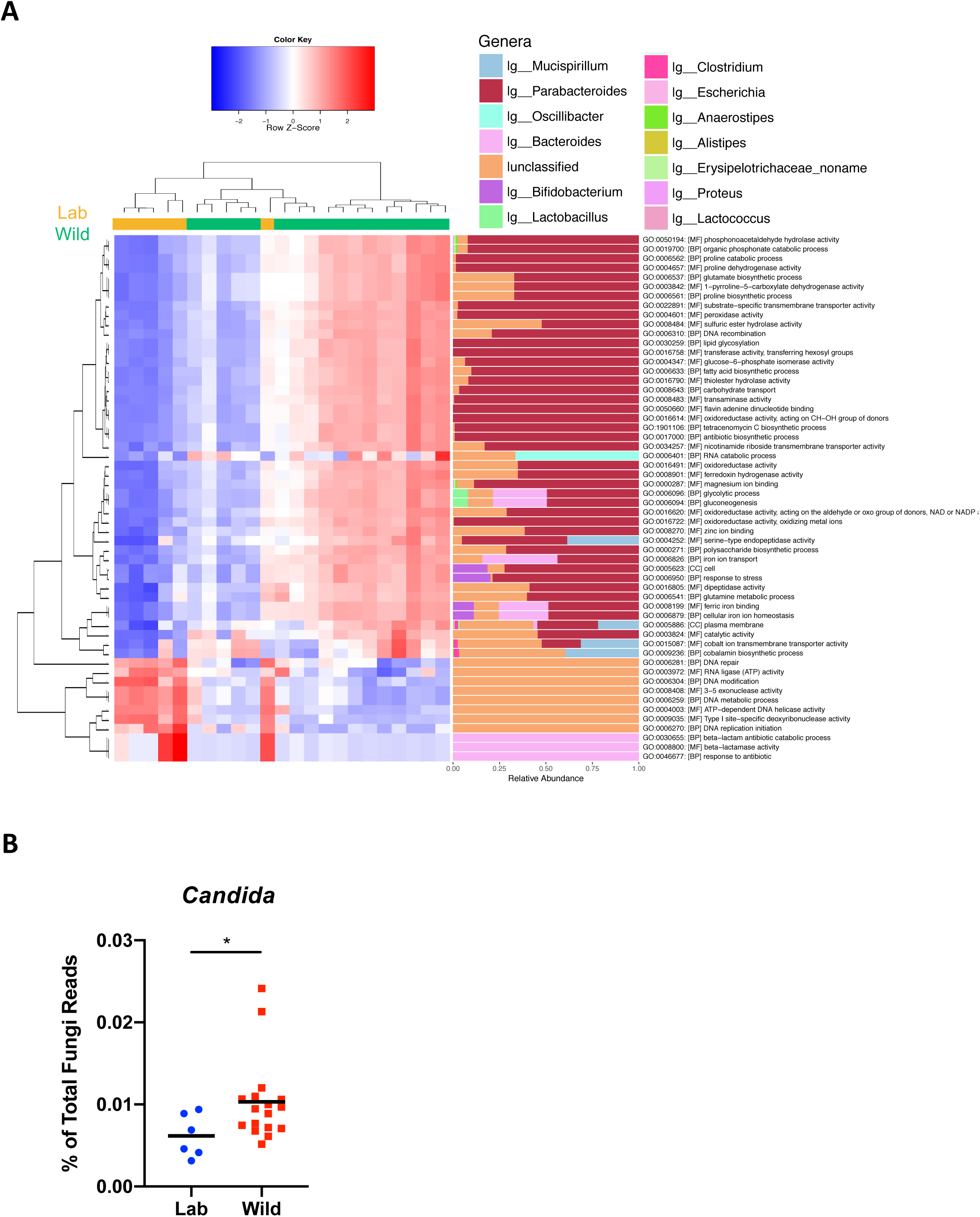
Rewilded mice have unique bacterial gene family signatures and are associated with an increase in *Candida* colonization. (**A**) Differential gene family abundance differences and changes in microbial genera taxa of lab versus rewilded mice. Differentially abundant gene families were evaluated by a fold change difference of 1.8 in either direction and an adjusted p-value of 0.05 from Student’s *t*-test between groups. (**B**) *Candida* specific reads identified in the shotgun sequencing. N = 6 lab and 17 rewilded mice.

**Supplemental Figure 4.**
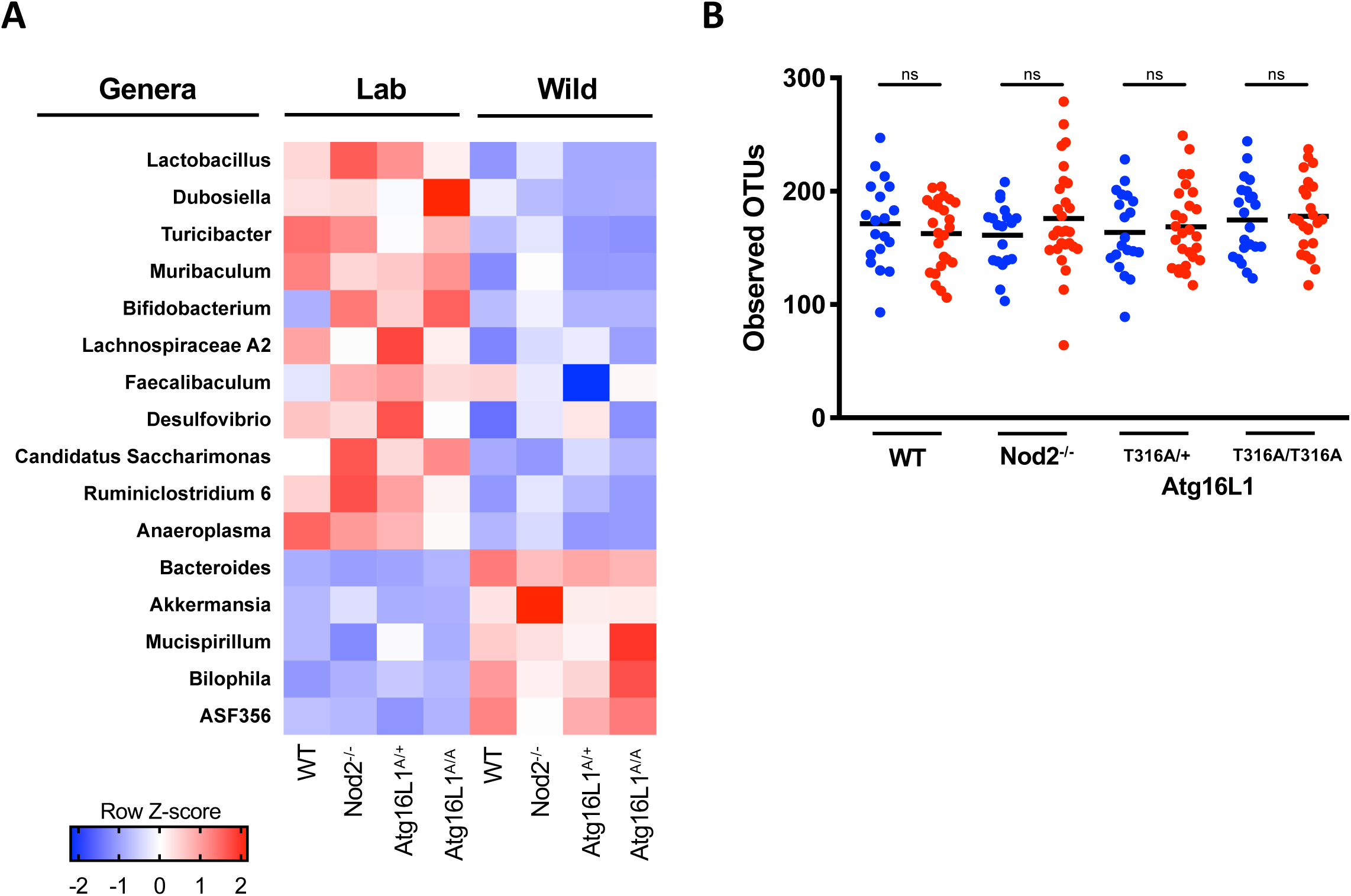
Bacterial community structure of wildtype, *Atg16l1*, and *Nod2* mutant mice in lab and wild conditions. (**A**) Heatmap of genera bacterial taxa significantly enriched in wildtype, *Atg16l1*, and *Nod2* mutant mice in lab and wild conditions as determined LEfSe analysis (Fig. 4D) (**B**) Alpha diversity quantification through number of OTUs in the above mice.

**Supplemental Figure 5.**
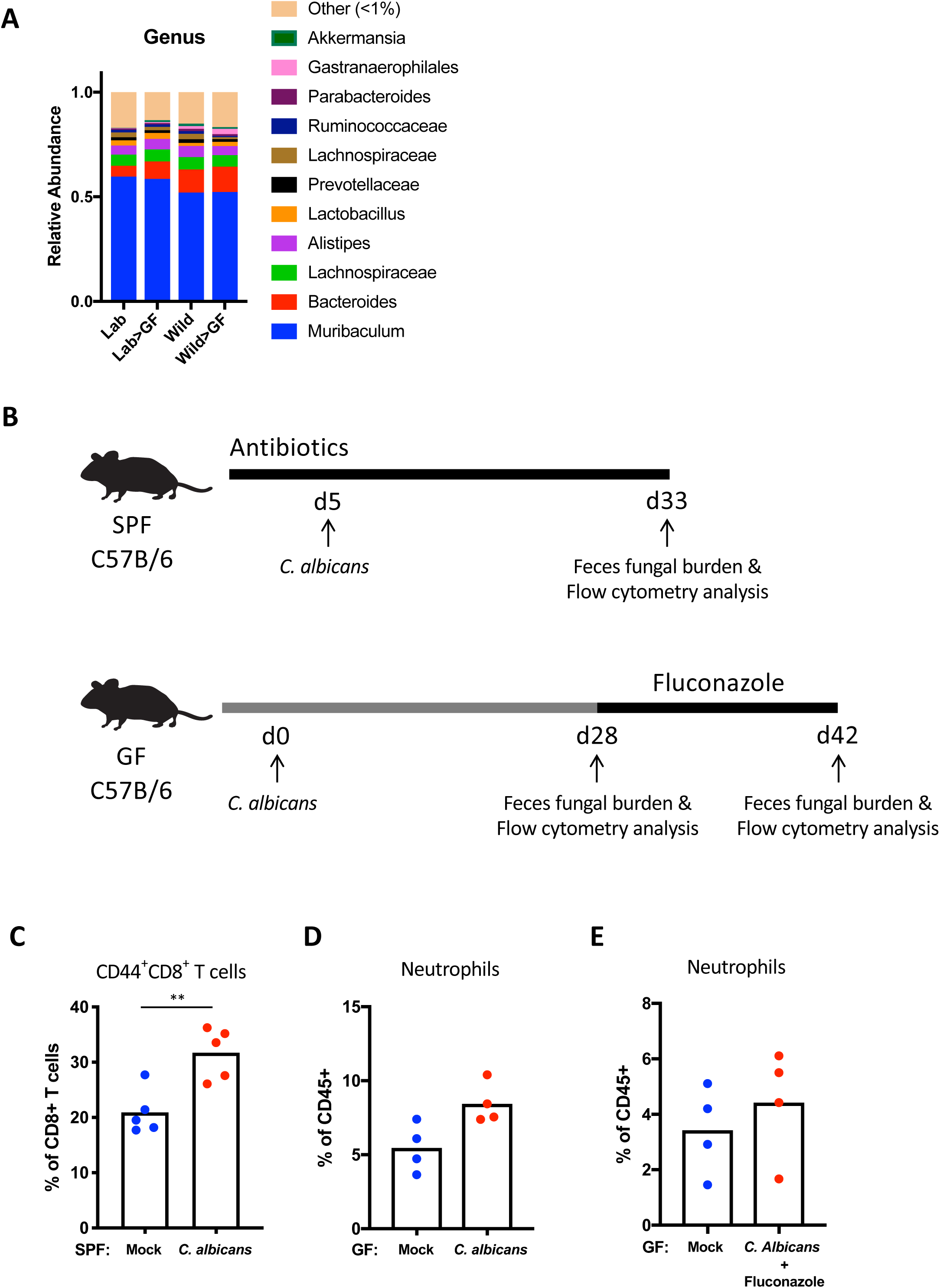
Bacterial community structure of germ-free mice reconstituted with microbiota from lab and rewilded mice. (**A**) Stacked bar plots displaying the mean relative abundances of genus level taxa within the fecal microbial communities in lab (Lab), rewilded (Wild) mice, or germ-free (GF) mice reconstituted with cecal contents from either lab or rewilded mice. N>5 recipient mice per condition. (**B**) Experimental model of *C. albicans* colonization in SPF and germ-free mice. (**C**) Quantification of CD44^+^CD8^+^ T cells in the peripheral blood of *C. albicans* colonized SPF mice. N = 5 mice per group. (**D**) Quantification of CD11b^+^Ly6G^+^ neutrophils in the peripheral blood of *C. albicans* mono-associated and control mice. N = 4 mice per group. (**E**) Quantification of neutrophils in the peripheral blood of *C. albicans* mono-associated mice after treatment with antifungal drug (Fluconazole) for 4 weeks. N = 4 mice per condition. ** *P* < 0.01 by two-tailed Student’s *t*-test between groups.

**Supplemental Figure 6.**
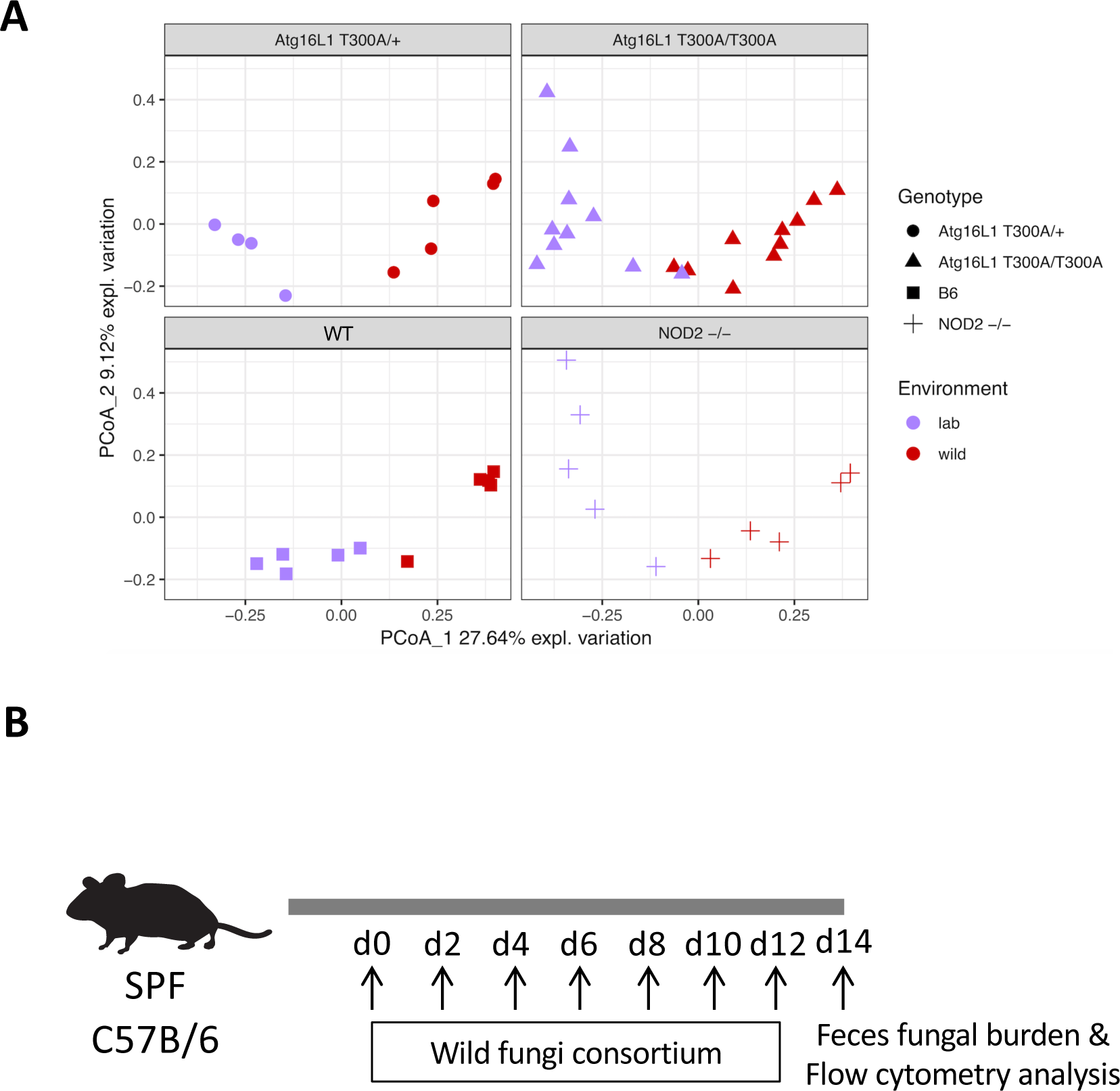
*Candida albicans* mono-colonization increases neutrophils in the peripheral blood. (**A**) PCA of mycobiota ITS sequences in wildtype, *Atg16l1*, and *Nod2* mutant mice in Lab and Wild conditions. (**B**) Experimental model of wild fungi colonization in SPF mice.

**Table 1.**
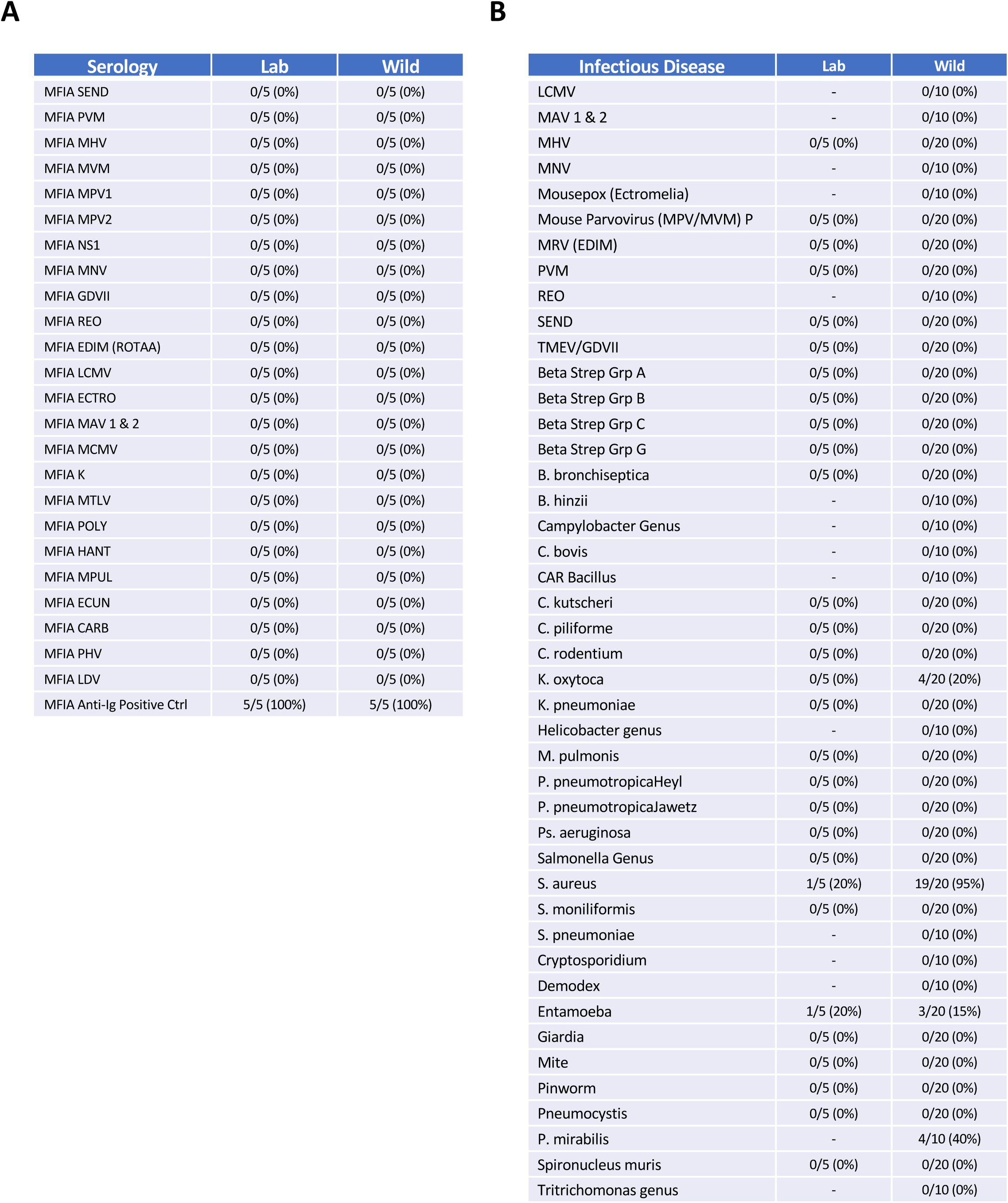
Rewilded mice are seronegative for common mouse pathogens and negative for most mouse pathogens and commensals. **(A)** Multiplexed Fluorometric ImmunoAssay results from serum of lab (Lab) and rewilded (Wild mice (Animal Health Diagnostic Services, MFIA Mouse Assessment Plus Profile, Charles River). N = 5 lab and 5 rewilded mice. (**B**) TaqMan PCR testing results from direct animal sampling of lab (Lab) and rewilded (Wild) mice (Animal Health Diagnostic Services, Surveillance Plus PRIA Panel, Charles River). N = 5 lab and 20 rewilded mice.

