## Supplementary material for "Altered immunity of laboratory mice in the natural environment is associated with fungal colonization": STAR METHODS

### STAR METODS

#### KEY RESOURCES TABLE

| REAGENT or RESOURCE | SOURCE | IDENTIFIER |
|---|---|---|
| Antibodies | | |
| Pacific Blue anti-mouse CD49b (pan-NK) Antibody | Biolegend | CAT#108918; RRID: AB_2265144 |
| Pacific Blue anti-mouse/human CD11b Antibody | Biolegend | CAT#101224; RRID: AB_755986 |
| Pacific Blue anti-mouse CD11c Antibody | Biolegend | CAT#117322; RRID: AB_755988 |
| Brilliant Violet 421 anti-mouse CD183 (CXCR3) Antibody | Biolegend | CAT#126529; RRID: AB_2563100 |
| Brilliant Violet 510 anti-mouse/rat/human CD27 Antibody | Biolegend | CAT#124229; RRID: AB_2565795 |
| Brilliant Violet 605 anti-mouse/human KLRG1 (MAFA) Antibody | Biolegend | CAT#138419; RRID: AB_2563357 |
| Brilliant Violet 785 anti-mouse CD3ε Antibody | Biolegend | CAT#100355 |
| Brilliant Violet 711 anti-mouse CD127 (IL-7Rα) Antibody | Biolegend | CAT#135035; RRID: AB_2564577 |
| PerCP/Cyanine5.5 anti-mouse CD279 (PD-1) Antibody | Biolegend | CAT#109120; RRID: AB_2566641 |
| APC/Cyanine7 anti-mouse CD4 Antibody | Biolegend | CAT#100414; RRID: AB_312699 |
| PE/Dazzle594 anti-mouse CD19 Antibody | Biolegend | CAT#115554; RRID: AB_2564001 |
| Brilliant Violet 650 anti-mouse CD8a Antibody | Biolegend | CAT#100742; RRID: AB_2563056 |
| Alexa Fluor 488 anti-mouse CD43 Activation-Associated Glycoform Antibody | Biolegend | CAT#121210; RRID: AB_528801 |
| APC anti-mouse CD62L Antibody | Biolegend | CAT#104412; RRID: AB_313099 |
| PE anti-mouse/human CD44 Antibody | Biolegend | CAT#103008; RRID: AB_312959 |
| Alexa Fluor 700 anti-mouse CD69 Antibody | Biolegend | CAT#104539; RRID: AB_2566304 |
| BUV395 Rat Anti-Mouse CD45 Antibody | BD Bioscience | CAT#564279; RRID: AB_2651134 |
| CD25 Monoclonal Antibody (PC61.5), PE-Cyanine7 | eBioscience | CAT#25-0251-82; RRID: AB_469608 |
| Pacific Blue anti-mouse/human CD45R/B220 Antibody | Biolegend | CAT#103227; RRID: AB_2566304 |
| Brilliant Violet 510 anti-mouse CD86 Antibody | Biolegend | CAT#105040; RRID: AB_2315766 |
| Brilliant Violet 605 anti-mouse CD3 Antibody | Biolegend | CAT#100237; RRID: AB_2562039 |
| Brilliant Violet 785 anti-mouse CD69 Antibody | Biolegend | CAT#104543; RRID: AB_2629640 |
| Alexa Fluor 488 anti-mouse CD40 Antibody | Biolegend | CAT#102910; RRID: AB_492852 |
| PerCP/Cyanine5.5 anti-mouse Ly-6G Antibody | Biolegend | CAT#127616; RRID: AB_1877271 |
| APC anti-mouse CD274 (B7-DC, PDL2) Antibody | Biolegend | CAT#127210; RRID: AB_2566345 |

| | | |
|---|---|---|
| APC/Cyanine7 anti-mouse IA/IE Antibody | Biolegend | CAT#107628; RRID: AB_2069377 |
| PE anti-mouse CD274 (B7-H1, PD-L1) Antibody | Biolegend | CAT#124308; RRID: AB_2073556 |
| PE/Dazzle594 anti-mouse CD64 (FCγRI) Antibody | Biolegend | CAT#139320; RRID: AB_2566559 |
| Alexa Fluor 700 anti-mouse F4/80 Antibody | Biolegend | CAT#123130; RRID: AB_2293450 |
| Brilliant Violet 650 anti-mouse CD11c Antibody | Biolegend | CAT#117339; RRID: AB_2562414 |
| BV421 Rat Anti-Mouse Siglec-F Antibody | BD Bioscience | CAT#562681; RRID: AB_2722581 |
| BV711 Rat Anti-Mouse CD103 Antibody | BD Bioscience | CAT#564320; RRID: AB_2738743 |
| PE-Cy7 Rat Anti-Mouse Ly-6C Antibody | BD Bioscience | CAT#560593; RRID: AB_1727557 |
| BUV395 Rat Anti-CD11b Antibody | BD Bioscience | CAT#563553; RRID: AB_2738276 |
| Bacterial and Fungal Strains | | |
| *Staphylococcus aureus* | K. Maurer *et al.,* 2015 | USA300 |
| *Pseudomonas aeruginosa* (PAO1) | D. Srivastava *et al.*, 2018 | PAO1 |
| *Bacillus subtilis* | ATCC | ATCC 6633 |
| *Clostridium perfringens* | NCNC | NCTC 10240 |
| *Bacteroides vulgatus* | ATCC | ATCC 8482 |
| *Candida albicans* | Dr. Stefan Feske, NYU | UC820 |
| *Candida albicans* | ATCC | SC-5314 |
| Chemicals, Peptides, and Recombinant Proteins | | |
| RBC lysis buffer | SANTA CRUZ | Cat#sc-296258 |
| HBSS | Gibco | Cat#14175-095 |
| BSA | Sigma | Cat#A7030 |
| EDTA | Invitrogen | Cat#15575-038 |
| HEPES | Corning | Cat#25-060-CI |
| Sodium pyruvate | Corning | Cat#25-000-CI |
| Critical Commercial Assays | | |
| Live/Dead Fixable Dead Cell Stain Kits | Invitrogen | Cat#L23105 |
| Custom mouse LEGENDplex assay | Biolegend | |
| NucleoSpin Soil Kit | Macherey-Nagel | Cat#740780.250 |
| RNeasy Plus Mini Kit | QIAGEN | Cat#74136 |
| Deposited Data | | |
| 16*S*, ITS, and RNA sequencing reads | NCBI Sequence Read Archive | PRJNA559026 |
| RNA expression counts | Gene Expression Omnibus | GSE135472 |

| | | |
|---|---|---|
| Cytokine and flow cytometry profiles | Github | https://github.com/ruggleslab/RewildedMice |
| Experimental Models: Organisms/Strains | | |
| Mouse: C57BL/6J | The Jackson Laboratory | JAX: 000664 |
| Mouse: *Nod2*$^{-/-}$ | Ramanan et. al., 2014 | |
| Mouse: *Atg16l1*$^{T316A/+}$ | Y. Matsuzawa-Ishimoto *et al.*, 2017 | |
| Mouse: *Atg16l1*$^{T316A/T316A}$ | Y. Matsuzawa-Ishimoto *et al.*, 2017 | |
| Software and Algorithms | | |
| Flowjo 10.4.2 | Flowjo, LLC | https://www.flowjo.com/ |
| Illustrator CC | Adobe | https://www.adobe.com/products/illustrator.html |
| Algorithms: t-SNE | fitsne v1.0.1 package | https://arxiv.org/abs/1712.09005 |
| Algorithms: UMAP | umap-learn v0.3.7 package | https://arxiv.org/abs/1802.03426 |
| Software: Python v3.6.5 | Python.org | https://www.python.org/downloads/release/python-365/ |
| Software: R v3.4.1 | | https://www.r-statistics.com/ |
| Algorithms: principal component analysis | ape v5.2 package | https://www.springer.com/gp/book/9781461417422 |
| Algorithms: Effect size measures | MDMR v0.5.1 package | https://link.springer.com/article/10.1007/s11336-016-9527-8 |
| Algorithms: random forest model | caret v6.0-80 package | R v3.4.1 |
| QIIME2 | Github | https://doc.qiime2.org |

#### LEAD CONTACT AND MATERIALS AVAILABILITY

#### EXPERIMENTAL MODEL AND SUBJECT DETAILS

**Mice and wild enclosure.**

All mouse lines were bred onsite in an MNV/Helicobacter-free specific pathogen free (SPF) facility at NYU School of Medicine to generate littermates from multiple breeding pairs that were randomly assigned to either remain in the institutional vivarium (lab mice) or released into the outdoor enclosures (rewilded mice) to control for the microbiota at the onset of the experiment. *Nod2*$^{-/-}$ and *Atg16l1*$^{T316A/T316A}$ mice on the C57BL/6J background were previously described (Matsuzawa-Ishimoto et al., 2017; Ramanan et al., 2014). *Atg16l1*$^{T316A/T316A}$ mice, *Atg16l1*$^{T316A/+}$, and wild-type (WT) control mice were generated from *Atg16l1*$^{T316A/+}$ breeder pairs, and *Nod2*$^{-/-}$ mice were generated from *Nod2*$^{-/-}$ breeder pairs. Additional C57BL/6J mice were purchased from Jackson Laboratory and bred onsite to supplement WT controls for experiments. 16S microbial diversity at the conclusion of the experiment did not show appreciable differences in microbial composition within the lab populations (see companion manuscript (Lin *et al*. Figure S5). Outdoor enclosures were previously described (Budischak et al., 2018; Leung et al., 2018) and the protocols for releasing the laboratory mice into the outdoor enclosure facility were approved by Princeton IACUC.

The enclosures consist of replicate outdoor pens, each measuring about 180 $m^2$ and fenced by 1.5-m high, zinced iron walls that are buried >80 cm deep and topped with electrical fencing to keep out terrestrial predators. Aluminum pie plates are strung up to deter aerial predators. A (180 × 140 × 70 cm) straw-filled shed is provided in each enclosure, along with two watering stations and a feeding station, so that the same mouse chow used in the laboratory (PicoLab Rodent Diet 20) was provided ad libitum to all mice. Mice outdoors, however, also had access to food sources found within the enclosures, including berries, seeds, and insects. 26-30 mice of mixed genotypes but the same sex were housed in each enclosure for 6-7 weeks. Longworth traps baited with chow were used to catch mice approximately 2 weeks and 4 weeks after release and again 6-7 weeks after release; for each trapping session, two baited traps were set per mouse per enclosure in the early evening, and all traps were checked within 12 hours. For subsequent microbiome assessment, a fresh stool sample was collected directly from the caught mice, flash frozen on dry ice, and stored at −80°C until further analysis. Mice were weighed with a spring balance.

30 WT, 29 *Nod2*$^{-/-}$, 31 *Atg16l1*$^{T316A/+}$, and 26 *Atg16l1*$^{T316A/T316A}$ laboratory mice (Total=116) were released into the outdoor enclosure. 19 WT, 19 *Nod2*$^{-/-}$, 20 *Atg16l1*$^{T316A/+}$, 22 *Atg16l1*$^{T316A/T316A}$ matched littermates (Total=80) were maintained in the institutional vivarium for comparison. For rewilded mice, traps were set regularly until the remaining mice were caught and were sampled for fecal microbiota. 25 WT, 28 *Nod2*$^{-/-}$, 27 *Atg16l1*$^{T316A/+}$, and 24 *Atg16l1*$^{T316A/T316A}$ rewilded mice (Total=104) were caught in the final trapping for terminal analyses. All lab control mice were recovered. Euthanasia was performed by CO2 asphyxiation, and blood, MLNs, and intestinal tissue were harvested. Two *Atg16l1*$^{T316A/+}$ rewilded mice failed quality control and were not included in downstream analyses. One *Atg16l1*$^{T316A/+}$ lab mouse was not appropriately processed and excluded in the final meta data table (Table S2; N=79 in lab mice and N=102 in rewilded mice).

#### METHOD DETAILS

##### Flow cytometry analysis.

At harvesting, MLNs were removed and the single-cell suspensions were prepared in FACS buffer (HBSS containing 1% BSA, 1mM EDTA, 20mM HEPES, and 1mM sodium pyruvate). The whole blood were also collected in a heparin containing tube and after centrifuging at 2000 rpm for 5 minutes, the designated plasma from supernatant was removed and stored at −80°C until all samples were collected and analyzed together. After two rounds of red blood cell lysis with 1x RBC lysis buffer for 5 minutes and wash with FACS buffer, the single-cell suspensions of whole blood cells were ready for the following staining procedure. MLN and whole blood cells were stained for live/dead with blue reactive dye and cell surface markers were labeled with the following antibody panels: Lymphoid panel: CD49b Pacific Blue, CD11b Pacific Blue, CD11c Pacific Blue, CXCR3 Brilliant Violet 421, CD27 Brilliant Violet 510, KLRG1 Brilliant Violet 605, CD3 Brilliant Violet 786, CD127 Brilliant Violet 711, PD1 PerCP/Cy5.5, CD4 APC/Cy7, CD19 PE/Dazzle594, CD8 Brilliant Violet 650, CD43 Alexa Fluor 488, CD62L APC, CD44 PE, CD69 Alexa Fluor 700, CD45 Buv395, CD25 PE/Cy7. Myeloid panel: B220 Pacific Blue, CD86 Brilliant Violet 510, CD3 Brilliant Violet 605, CD69 Brilliant Violet 786, CD40 Alexa Fluor 488, Ly6G PerCP/Cy5.5, PDL2 APC, IA/IE APC/Cy7, PDL1 PE, CD64 PE/Dazzle594, F4/80 Alexa Fluor 700, CD11c Brilliant Violet 650, Siglec-F Brilliant Violet 421, CD103 Brilliant Violet 711, Ly6C PE/Cy7, CD11b Buv395. FACS analyses were performed in a ZE5 cell analyzer (BIO-RAD) and recorded FACS data were analyzed by Flowjo v10.4.2.

##### MLN cell stimulation and cytokine profiling

Single cell suspension of MLN cells were reconstituted in RPMI at $2 \times 10^6$ cells/mL, and 0.1 mL was cultured in 96-well microtiter plates that contained $10^7$ cfu/mL UV-killed microbes, $10^5$ αCD3/CD28 beads, or PBS control. Overnight microbial cultures were reconstituted at $10^8$ cfu/mL prior to irradiation. The stimulated microbes are as following: *Staphylococcus aureus* (Maurer et al., 2015), *Pseudomonas aeruginosa* (PAO1) (kindly provided by Dr. Andrew Darwin, NYU) (Srivastava et al., 2018), *Bacillus subtilis* (ATCC 6633), *Clostridium perfringens* (NCTC 10240), *Bacteroides vulgatus* (ATCC 8482), and *Candida albicans* (UC820, kindly provided by Dr. Stefan Feske, NYU). Supernatants were collected after 2 days and stored at −80°C. Concentrations of IL-1α, IL-1β, IL-4, IL-5, IL-6, IL-10, IL-13, IL-17A, CCL2, CCL3, CCL4, CXCL1, IFN-γ, and TNF-α in supernatants were measured using a custom mouse LEGENDplex assay (Biolegend) according to the manufacturer's instructions. Plasma concentrations of IL-1α, IL-1β, IL-6, IL-10, RANTES, CCL2, CCL3, CCL4, CCL20, CXCL1, CXCL10, TNFα, GM-CSF were measured using a second custom mouse LEGENDplex assay (Biolegend). For each mouse profiled for cytokine production in response to microbial stimuli a PBS control was also sampled. In order to normalize per mouse cytokine production, we calculated the fold change of each cytokine measure to the PBS control. There was no overall difference in baseline cytokine production for any cytokine in response to PBS between lab and rewilded mice.

**16S library preparation and sequencing**

DNA was isolated from stool samples using the NucleoSpin Soil Kit (Macherey-Nagel). Bacterial 16S rRNA gene was amplified at the V4 region using primer pairs and paired-end amplicon sequencing was performed on the Illumina MiSeq system as previously described (Neil et al., 2019). Sequencing reads were processed using the DADA2 pipeline in the QIIME2 software package. Taxonomic assignment was performed against the Silva v132 database. Alpha diversity analysis was done using observed OTUs. Differential abundance taxa were identified using linear discriminant analysis effect size (LefSE) in different biological groups at a threshold LDA score described in the legends (Segata et al., 2011).

**Whole shotgun metagenomic sequencing**

Shotgun metagenomics sequence data was analyzed from rewilded and lab mice for microbial abundance and gene family abundance profiles as previously described (Devlin et al., 2018). In brief, the sequencing data were processed through an analysis pipeline utilizing the Huttenhower Biobakery pipeline (McIver et al., 2018), including FastQC, Kneaddata, MetaPhlAn (Truong et al., 2015) and HUMAnN2 (Abubucker et al., 2012) to obtain an annotated gene abundance matrix. In order to discern major differences between lab and rewilded mice we utilized the WHAM! (Devlin et al., 2018) interactive analysis platform and subsequent statistical tests to uncover major gene family abundance differences and changes in microbial taxa. Differentially abundant gene families were evaluated by a fold change difference of 1.8 in either direction of lab versus rewilded mice and an adjusted p-value of 0.05 from a t-test. The abundance of these differentially significant gene families was also used for principal component analysis.

**Pathogen screen**

Randomly selected mice were screened for the presence of infectious agents using EZ-Spot Assessment Plus Multiplexed Fluorometric Immunoassay (MFIA) and PCR Infectious Agent Testing (PRIA) Surveillance Plus Panel (Charles River Laboratories). Dried blood, feces, and body swabs were collected according to submission guidelines.

**MLN cell RNA preparation and sequencing**

Frozen samples of single cell suspensions from MLN of lab or rewilded mice were thaw to isolate RNA from approximately $10^6$ cells by RNeasy Plus Mini Kit according to manufacturer's instructions. CEL-seq2 were performed to do RNA sequencing on samples with good RNA qualities (RNA integrity number $\geqq$ 5)

**Histology and imaging**

Immunohistochemistry was performed on 10% neutral buffered formalin fixed, paraffin-embedded small intestine and colon tissue. Sections were collected at 5-microns onto plus slides (Fisher Scientific, Cat #

22-042-924) and stored at room temperature prior to use. Unconjugated, rabbit anti-mouse CD 8a, T-cell surface glycoprotein alpha chain, clone D4W2Z (Cell Signaling Cat# 98941 Lot# 1 RRID: unassigned) raised against synthetic peptide corresponding to residues surrounding Asp42 of mouse CD8α protein (Russell et al., 2018) was used for immunohistochemistry.

**Transfer of wild microbiota**

Flash frozen cecal contents from rewilded mice were transferred by two consecutive daily oral gavage procedures into timed pregnant germ-free mice as previously described (Rosshart et al., 2017). Three representative lab and rewilded wildtype mice (defined as within a standard deviation of the average number of blood granulocytes in the respective conditions) served as donors of cecal contents. Each recipient germ-free dam received the same 0.1 to 0.15 mL suspension on day 14 and 15 of pregnancy. 16S rRNA sequencing was performed as described above.

**Wild enterobacteriaceae isolation**

Serial dilutions of cecal contents from rewilded or lab mice were plated on MacConkey Agar and incubated overnight at 37$^{o}$C. Colonies grew from rewilded samples within 12–16 hours but not lab samples. Colonies were chosen at random and were identified by 16S rRNA sequencing. For inoculation into mice, unique species was cultured in LB broth at 37°C for 16 hrs, pooled, and conventional mice were gavaged with $1.5\times10^{8}$ bacteria. Stool was collected after two weeks and bacterial burden was determined.

**Staphylococcus aureus isolation**

Fecal bacteria were quantified by dilution plating on CHROMagar Staph aureus plates at 37°C. Colonies were selected at random and genotyped by DNA sequence analysis of the protein A gene variable repeat region (spa typing) and a variety of additional DNA polymorphisms and the presence of pvl genes as previously described (Copin et al., 2019).

**Fecal Fungi quantification by qPCR**

Fungal DNA was isolated from individual fecal pellets using the NucleoSpin Soil kit (Macherey-Nagel) as described in 16S library preparation. Quantitative PCR was performed using SybrGreen (Roche) on a Roche480II Lightcycler using the following primers: Fungal ITS1-2, Fwd 5'-CTTGGTCATTTAGAG-GAAGTAA-3' and Rev 5'-GCTGCGTTCTTCATCGATGC-3'. Relative abundance of fungal-specific internal transcribed spacer (ITS) rDNA was calculated using the $\Delta C_T$ method and the values were converted as fold change from the average $C_T$ of control lab mice samples.

**ITS library preparation and sequencing**

The fungal ITS2 region was targeted for amplification and sequencing with primers that append barcodes and Illumina adapters, similar to those described by (Taylor et al., 2016). The full primer sequences used were (with XXXXXX indicating barcode sequences, and bold lettering for the ITS2 targeting primers): AATGATACGGCGACCACCGAGATCTACACXXXXXXXXACACTCTTTCCCT ACACGACGCTCTTCCGATCTAAAGCCTCCGCTTATTGATATGCTTAART, CAAGCAGAAGACGGCATACGAGATXXXXXXXXGTGACTGGAGTTCAGAC GTGTGCTCTTCCGATCTGGAACTTTYRRCAAYGGATCWCT. Notably, this results in the Illumina read-one from the reverse (LSU) side of the ITS reference sequence and the read-two from the 5.8S side. The ITS2 region was amplified and Illumina adapters appended by PCR in 25 µl volume with Q5 High-Fidelity 2X master mix (NEB, # M0492L). PCR conditions were as follows: initial denaturation at 95°C for 2 minutes, followed by 32 cycles of 98°C for 15 seconds, 52°C for 20 seconds, 72°C for 45 seconds, and a final 2 minute extension at 72°C. We performed triplicate PCR reactions for each sample, then combined triplicates and cleaned the PCR products with Axygen AxyPrep Mag PCR cleanup beads (Corning, #MAG-PCR-CL-50) using 1.8X volumes of beads diluted to 62.5% in water (v/v) to remove large primer-dimers. Cleaned products were quantified then multiplexed at even abundances and sequenced on an Illumina MiSeq in paired-end mode with 300 cycles each and a 5% PhiX library spike-in to provide increased base diversity. All sequences have been deposited at NCBI under the BioProject PRJNA559026.

**ITS sequences analysis**

Raw sequences were imported into QIIME2 and read-pairs were joined with the vsearch plugin requiring a minimum length of 300 nucleotides (a completely overlapping pair of reads). Joined reads were then quality filtered and exported to reverse-complement reads with the fastx toolkit (http://hannonlab.cshl.edu/fastx_toolkit/) due to the orientation of our ITS2 primers. Joined, quality-filtered and reverse-complemented reads were re-imported into QIIME2 to employ the ITSxpress plugin in order to trim primers and extract the ITS2 sub-region (Bengtsson-Palme et al., 2013; Rivers et al., 2018). ITS2 sub-region extracted sequences were then denoised with deblur and a trim-length of 150 against the UNITE fungal database reference sequences (version 7) (Amir et al., 2017; Nilsson et al., 2018). Finally, the UNITE database was used to train a naïve Bayes classifier and to assign fungal taxonomies in QIIME2 (Bokulich et al., 2018). The community matrix was subsampled at a depth of 906 sequences per sample and Bray-Curtis distances and alpha diversity metrics (observed otus) were calculated within QIIME2.

***Candida albicans* inoculation**

The *C. albicans* laboratory strain SC5314 was purchased from the ATCC. Fungi were cultured in Sabouraud Dextrose media with chloramphenicol (25 µg/ml, Sigma) at 30$^{o}$C for 16 hours. The culture then was washed and suspended in PBS. Antibiotic treated or germ-free mice were orally gavaged with 150 µl of *C. albicans* ($10^{7}$ fungal CFUs). Feces and organs were collected and analyzed after 4 weeks colonization. For depletion of colonized *C. albicans*, germ-free mice were supplemented with fluconazole (0.5 mg/ml) in drinking water for another 2 weeks. Feces and organs were collected accordingly for analysis.

**Isolation of Wild fungi**

Feces or ileocecal contents from re-wilded mice were resuspended in PBS and plated on Sabouraud Dextrose Agar (SDA) with chloramphenicol (25 µg/ml, Sigma) at 25 $^{o}$C for 3 days. Individual colonies were re-grown on SDA plates to acquire the pure strains. Fungal DNA then was extracted from each isolated strains and sequenced using the ITS primers described below to identify the fungal species.

**Wild fungal consortium inoculation**

A wild fungi consortium consisting of *A. candidus*, 4 strains of *A. proliferans*, *C. globosum* and *D. Indicus* (corresponding to the fungi on plates (i)–(vii) from Figure 4E was cultured on SDA for 5 days and further cultured in Sabouraud Dextrose broth at 25$^{o}$C for 2 days. Fungi were passed through a 19-gauge syringe needle several times to break down the mycelium prior to oral gavage into mice and mixed at equal ratios. SPF mice were orally gavaged with 150 µl of wild fungi consortium every other day for 2 weeks and germ-free mice were giving with 150 µl of wild fungi consortium at the first day of experiment.

**Statistical analysis**

An unpaired two-tailed t test was used to evaluate differences between two groups. An ANOVA was used to evaluate experiments involving multiple groups with the Holm-Sidak multiple comparisons test.

#### DATA AND CODE AVAILABILITY

Raw sequence data from 16*S*, ITS, and RNA sequencing experiments are deposited in the NCBI Sequence Read Archive under BioProject accession number PRJNA559026 and gene expression omnibus (GEO) accession number GSE135472. All processing was performed in R and analysis scripts can be found on Github at https://github.com/ruggleslab/RewildedMice

#### METHOD REFERENCE

**Abubucker, S., Segata, N., Goll, J., Schubert, A.M., Izard, J., Cantarel, B.L., Rodriguez-Mueller, B., Zucker, J., Thiagarajan, M., Henrissat, B., White, O., et al. (2012) Metabolic reconstruction for metagenomic data and its application to the human microbiome. PLoS Comput. Biol. 8(6), e1002358..**

**Amir, A., McDonald, D., Navas-Molina, J.A., Kopylova, E., Morton, J.T., Zech Xu, Z., Kightley, E.P., Thompson, L.R., Hyde, E.R., Gonzalez, A., et al. (2017) Deblur Rapidly Resolves Single-Nucleotide Community Sequence Patterns. mSystems. 2(2), e00191-16.**

**Bengtsson-Palme, J., Ryberg, M., Hartmann, M., Branco, S., Wang, Z., Godhe, A., De Wit, P., Sánchez-García, M., Ebersberger, I., de Sousa, F., et al. (2013). Improved software detection and extraction of ITS1 and ITS2 from ribosomal ITS sequences of fungi and other eukaryotes for analysis of environmental sequencing data. Methods in Ecology and Evolution. 4(10), 914-919. DOI: 10.1111/2041-210X.12073.**

**Bokulich, N.A., Kaehler, B.D., Rideout, J.R., Dillon, M., Bolyen, E., Knight, R., Huttley, G.A., and Gregory Caporaso, J. (2018) Optimizing taxonomic classification of marker-gene amplicon sequences with QIIME 2's q2-feature-classifier plugin. Microbiome. 6(1), 90.**

**Budischak, S.A., Hansen, C.B., Caudron, Q., Garnier, R., Kartzinel, T.R., Pelczer, I., Cressler, C.E., van Leeuwen, A., and Graham, A.L. (2018). Feeding Immunity: Physiological and Behavioral Responses to Infection and Resource Limitation. Front Immunol. 8, 1914. Published online 2018/01/24 DOI: 10.3389/fimmu.2017.01914.**

**Copin, R., Sause, W.E., Fulmer, Y., Balasubramanian, D., Dyzenhaus, S., Ahmed, J.M., Kumar, K., Lees, J., Stachel, A., Fisher, J.C., et al. (2019). Sequential evolution of virulence and resistance during clonal spread of community-acquired methicillin-resistant Staphylococcus aureus. Proceedings of the National Academy of Sciences of the United States of America. 116(5), 1745-1754. Published online 2019/01/13 DOI: 10.1073/pnas.1814265116.**

**Devlin, J.C., Battaglia, T., Blaser, M.J., and Ruggles, K.V. (2018) WHAM!: a web-based visualization suite for user-defined analysis of metagenomic shotgun sequencing data. BMC Genomics. 19(1), 493.**

**Leung, J.M., Budischak, S.A., Chung The, H., Hansen, C., Bowcutt, R., Neill, R., Shellman, M., Loke, P., and Graham, A.L. (2018). Rapid environmental effects on gut nematode susceptibility in rewilded mice. PLoS biology. 16(3), e2004108. Published online 2018/03/09 DOI: 10.1371/journal.pbio.2004108.**

**Matsuzawa-Ishimoto, Y., Shono, Y., Gomez, L.E., Hubbard-Lucey, V.M., Cammer, M., Neil, J., Dewan, M.Z., Lieberman, S.R., Lazrak, A., Marinis, J.M., et al. (2017). Autophagy protein ATG16L1 prevents necroptosis in the intestinal epithelium. J Exp Med. 214(12), 3687-3705. Published online 2017/11/02 DOI: jem.20170558 [pii] 10.1084/jem.20170558.**

**Maurer, K., Reyes-Robles, T., Alonzo, F., 3rd, Durbin, J., Torres, V.J., and Cadwell, K. (2015). Autophagy mediates tolerance to Staphylococcus aureus alpha-toxin. Cell Host Microbe. 17(4), 429-440. Published online 2015/03/31 DOI: S1931-3128(15)00116-X [pii] 10.1016/j.chom.2015.03.001.**

McIver, L.J., Abu-Ali, G., Franzosa, E.A., Schwager, R., Morgan, X.C., Waldron, L., Segata, N., and Huttenhower, C. (2018). bioBakery: a meta'omic analysis environment. Bioinformatics (Oxford, England). 34(7), 1235-1237. Published online 2017/12/02 DOI: 10.1093/bioinformatics/btx754.

Neil, J.A., Matsuzawa-Ishimoto, Y., Kernbauer-Holzl, E., Schuster, S.L., Sota, S., Venzon, M., Dallari, S., Galvao Neto, A., Hine, A., Hudesman, D., et al. (2019). IFN-I and IL-22 mediate protective effects of intestinal viral infection. Nat Microbiol. Published online 2019/06/12 DOI: 10.1038/s41564-019-0470-110.1038/s41564-019-0470-1 [pii].

Nilsson, R.H., Larsson, K.H., Taylor, A.F.S., Bengtsson-Palme, J., Jeppesen, T.S., Schigel, D., Kennedy, P., Picard, K., Glockner, F.O., Tedersoo, L., et al. (2019) The UNITE database for molecular identification of fungi: handling dark taxa and parallel taxonomic classifications. Nucleic Acids Res. 47(D1), D259-D264.

Ramanan, D., Tang, M.S., Bowcutt, R., Loke, P., and Cadwell, K. (2014). Bacterial sensor Nod2 prevents inflammation of the small intestine by restricting the expansion of the commensal Bacteroides vulgatus. Immunity. 41(2), 311-324. Published online 2014/08/05 DOI: S1074-7613(14)00241-6 [pii]10.1016/j.immuni.2014.06.015.

Rivers, A.R., Weber, K.C., Gardner, T.A., Liu, S., and Armstrong, S.D. (2018) ITSxpress: Software to rapidly trim internally transcribed spacer sequences with quality scores for marker gene analysis. F1000Res. 7, 1418.

Rosshart, S.P., Vassallo, B.G., Angeletti, D., Hutchinson, D.S., Morgan, A.P., Takeda, K., Hickman, H.D., McCulloch, J.A., Badger, J.H., Ajami, N.J., et al. (2017). Wild Mouse Gut Microbiota Promotes Host Fitness and Improves Disease Resistance. Cell. 171(5), 1015-1028.e1013. Published online 2017/10/24 DOI: 10.1016/j.cell.2017.09.016.

Russell, L., Swanner, J., Jaime-Ramirez, A.C., Wang, Y., Sprague, A., Banasavadi-Siddegowda, Y., Yoo, J.Y., Sizemore, G.M., Kladney, R., Zhang, J., et al. (2018) PTEN expression by an oncolytic herpesvirus directs T-cell mediated tumor clearance. Nat Commun. 9(1), 5006.

Srivastava, D., Seo, J., Rimal, B., Kim, S.J., Zhen, S., and Darwin, A.J. (2018). A Proteolytic Complex Targets Multiple Cell Wall Hydrolases in Pseudomonas aeruginosa. MBio. 9(4). Published online 2018/07/19 DOI: mBio.00972-18 [pii]10.1128/mBio.00972-18.

Taylor, D.L., Walters, W.A., Lennon, N.J., Bochicchio, J., Krohn, A., Caporaso, J.G., and Pennanen, T. (2016) Accurate Estimation of Fungal Diversity and Abundance through Improved Lineage-Specific Primers Optimized for Illumina Amplicon Sequencing. Appl Environ Microbiol. 82(24), 7217-7226.

Truong, D.T., Franzosa, E.A., Tickle, T.L., Scholz, M., Weingart, G., Pasolli, E., Tett, A., Huttenhower, C., and Segata, N. (2015) MetaPhlAn2 for enhanced metagenomic taxonomic profiling. Nat Methods. 12(10), 902-903.
